# Subthreshold serotonin signals combined by the G proteins Gα_q_ and Gα_s_ activate the *C. elegans* egg-laying muscles

**DOI:** 10.1101/2022.09.12.507489

**Authors:** Andrew C. Olson, Allison M. Butt, Nakeirah T.M. Christie, Ashish Shelar, Michael R. Koelle

## Abstract

Individual neuron or muscle cells express many G protein coupled receptors (GPCRs) for neurotransmitters and neuropeptides. It remains unclear how these cells integrate multiple GPCR signals that all must act through the same few G proteins. We investigated how two serotonin GPCRs, Gα_q_-coupled SER-1 and Gα_s_-coupled SER-7, function together on the *C. elegans* egg-laying muscles to promote contraction and thus cause eggs to be laid. Using receptor null mutations and cell-specific knockdowns, we found that serotonin signaling through either SER-1/Gα_q_ or SER-7/Gα_s_ alone does not induce egg laying, but these subthreshold signals can combine to promote egg laying. However, using designer receptors or optogenetics to artificially induce high levels of either Gα_q_ signaling or Gα_s_ signaling in the muscles was sufficient to induce egg laying. Conversely, knocking down both Gα_q_ and Gα_s_ in the egg-laying muscle cells induced egg-laying defects stronger than those of a *ser-7 ser-1* double knockout. These results suggest that, in the egg-laying muscles, multiple GPCRs for serotonin and other signals each produce weak effects that individually do not result in strong behavioral outcomes. However, they can combine to produce sufficient levels of Gα_q_ and Gα_s_ signaling to promote muscle activity and egg laying.

## Introduction

Individual neuron or muscle cells can express many different G protein coupled receptors (GPCRs), which in turn act through just three main types of heterotrimeric G proteins: G_s_, G_q/11_, and G_i/o_ (Kaur et al., 2017; Smith et al., 2019; Jiang et al., 2022). Evidently, many different chemical signals impinge on an individual cell within the body, and signaling through multiple GPCRs integrates this complex information to produce appropriate responses. How this occurs remains largely unclear and is the focus of this study.

Evidence for the widespread use of multiple GPCRs on individual cells to integrate chemical signals comes from studies across different cell types and organisms. Single-cell transcriptomics on primary cultures of mouse smooth muscle cells and endothelial cells indicate that individual cells express ∼20 GPCRs on average (Kaur et al., 2017). Even when only 29 out of the >100 neuropeptide receptor genes are analyzed, a typical neuron expresses multiple such receptors (Smith et al., 2019). Vertebrate mast cells use at least 16 different GPCRs to respond to various neurotransmitters and neuropeptides (Xu et al., 2020).

Neural circuits of invertebrates that consist of only a small number of cells provide model systems in which it is possible to tease out how multiple GPCRs function together on individual cells. For example, in the crustacean somatogastric circuit, indirect evidence suggests a large number of different neurotransmitters and neuropeptides modulate activity of individual neurons (Marder and Bucher, 2007). In this study, we focus on the *C. elegans* egg-laying circuit, where we recently found that individual neuron and muscle cells each express multiple neurotransmitter GPCRs (Fernandez et al., 2020). A pair of neurons in this circuit release serotonin to activate egg laying, and there are multiple different serotonin GPCRs co-expressed on individual cells in the circuit. Thus, serotonin signaling in the *C. elegans* egg-laying circuit provides an opportunity to analyze how different GPCRs on the same cells function together.

There are multiple serotonin GPCRs in both humans and *C. elegans*, and these are often co-expressed on the same target cells (Feng et al., 2001; Bonn et al., 2013). Humans have six families of serotonin GPCRs comprising at least 12 receptor subtypes (Sarkar et al., 2021). Pyramidal neurons, for example, can express up to five different subtypes of serotonin receptors, including two different Gα_q_-coupled 5HT_2_ receptor subtypes and the Gα_s_-coupled 5HT_4_ receptor (Feng et al., 2001). Signaling through each of these Gα_s_- and Gα_q_-coupled serotonin receptors can increase the excitability of target neurons (Rasmussen and Aghajanian, 1990; Lopez et al., 2021), but the logic of using multiple serotonin receptors in parallel on the same target cells remains unclear.

In the *C. elegans* egg-laying circuit, schematized in Figure 1A, the hermaphrodite specific neurons (HSNs) and ventral type C (VC) motor neurons synapse onto the egg-laying muscles. The HSNs release both serotonin and a neuropeptide named NLP-3 to induce activity of the VCs and contraction of the egg-laying muscles, resulting in egg laying (Collins and Koelle, 2013; Brewer et al., 2019). There are 16 egg-laying muscle cells in total, four each of four types: the um1 and um2 uterine muscle cell types, as well as vm1 and vm2 vulval muscle cell types. The um1, um2, and vm2 muscle cells each co-express the two serotonin receptors, SER-1 and SER-7, that are principally responsible for inducing egg laying (Fernandez et al., 2020). SER-1 is a Gα_q_-coupled receptor, while SER-7 couples to Gα_s_ (Hamdan et al., 1999; Hobson et al., 2003; Carnell et al., 2005; Dempsey et al., 2005; Carre-Pierrat et al., 2006; Hobson et al., 2006). The vm1 muscle cells as well as the VC4 and VC5 neurons each express SER-7 but not SER-1 (Fernandez et al., 2020).

**Figure 1.**
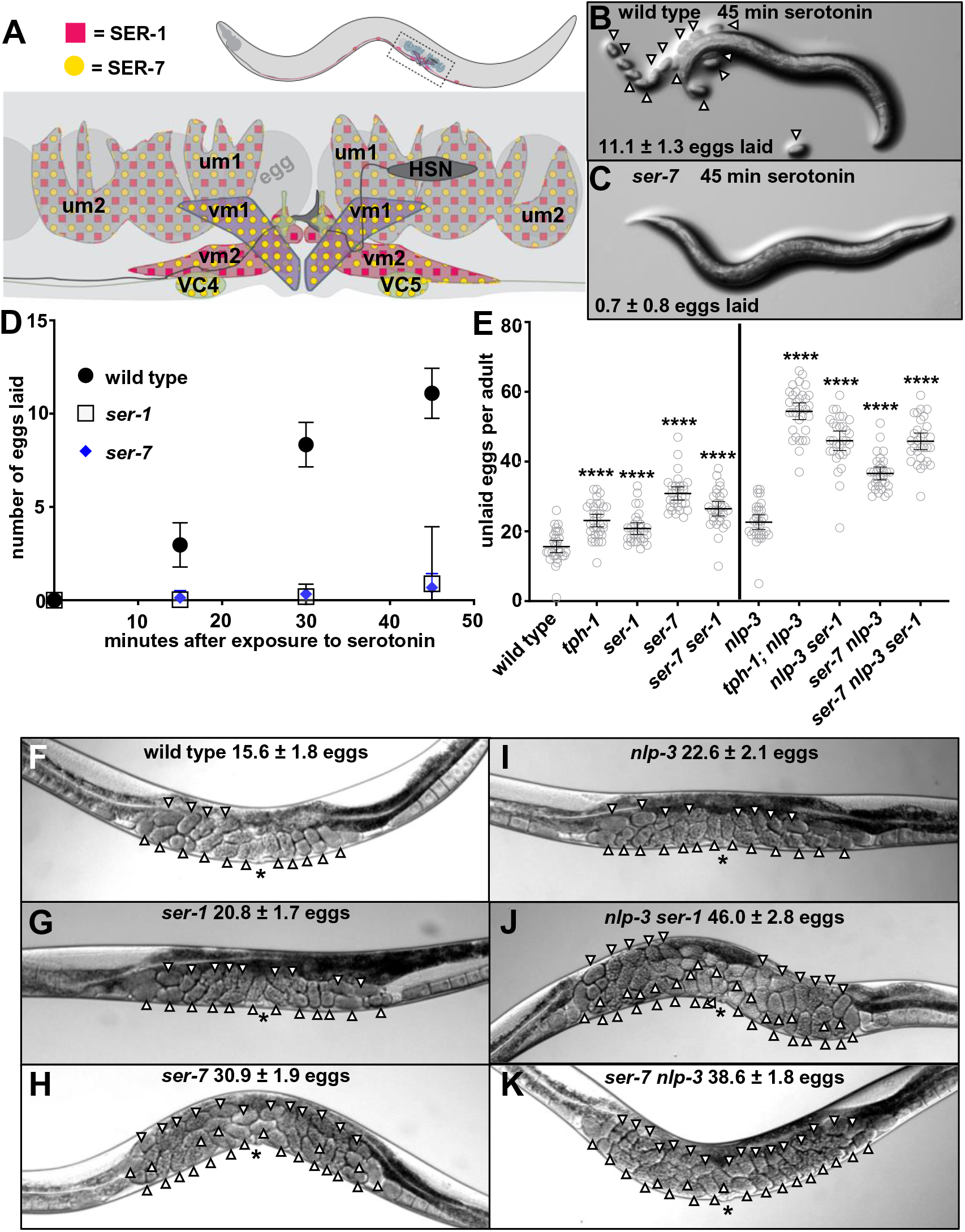
The serotonin receptors SER-1 and SER-7 are co-expressed on cells of the egg-laying circuit and loss of either blocks the ability of serotonin to stimulate egg laying. (**A**) Schematic of the *C. elegans* egg-laying system. Yellow circles denote cells that express SER-1 and pink squares denote cells that express SER-7. The HSN neurons and the vm1, vm2, um1, and um2 muscle cells each occur in left/right pairs, but only the cells on the left side of the animal are shown in this schematic. VC4 and VC5 are single neurons. (**B, C**) Serotonin-induced egg-laying assays for wild-type or *ser-7* knockout worms. Worms were photographed 45 minutes after being placed on plates containing 26 mM serotonin. Serotonin partially paralyzed the worms so that they remained adjacent to their laid eggs, which are indicated by arrowheads. The average number of laid eggs for each genotype in this assay is shown. (**D**) Results of a time course using the same assay illustrated in panels **B** and **C** for wild-type, *ser-1*, and *ser-7* null mutant animals. The assay was repeated with 10 worms/plate at least 3 times per genotype. (**E**) Average number of unlaid eggs per adult worm, n≥30 for each genotype. Genotypes left of the vertical black line are statistically compared to wild-type control animals. Genotypes right of the line are statistically compared to *nlp-3* single-mutant control animals. **** = p<0.0001 for these comparisons. (**F-K**) Photographs of individual worms illustrating the accumulation of unlaid eggs (indicated by white arrowheads) in some of the genotypes analyzed in (**E**). The vulval slit is indicated by *. Photographs of individual worms for the remaining genotypes are shown in Figure 1-figure supplement 1. The average number of unlaid eggs for each genotype is also indicated. All measurements are given with 95% confidence intervals.

In this study, we have analyzed how SER-1/Gα_q_ signaling and SER-7/Gα_s_ signaling act together in the egg-laying circuit to result in egg laying.

## Results

### Both the SER-1 and SER-7 serotonin receptors are required for exogenous and endogenous serotonin to stimulate egg laying

Application of exogenous serotonin causes wild-type worms to quickly initiate egg laying (Figure 1B and D). We reproduced previous studies (Carnell et al., 2005; Dempsey et al., 2005; Hobson et al., 2006) showing that SER-1 and SER-7 are each required for such serotonin-induced egg laying (Figure 1B-D). Animals carrying null alleles of the *ser-1* (Figure 1D) or *ser-7* (Figure 1C and D) genes each showed severely reduced egg laying in response to exogenous serotonin. While one might have expected these co-expressed receptors to function redundantly, resulting in weak defects when knocking out one or the other, the surprisingly strong defects seen in the single receptor mutants prompted us to analyze in depth how SER-1 and SER-7 function together.

A third serotonin receptor, SER-5, has been reported to weakly promote serotonin-induced egg laying in certain genetic backgrounds (Hapiak et al., 2009). Because this effect is so weak (Figure 1-figure supplement 1) and SER-5 expression in the egg-laying system is reported as either weak and variable (Hapiak et al., 2009) or undetectable (Fernandez et al., 2020), this study excludes SER-5 from further analysis.

To reveal how SER-1 and SER-7 receptors mediate the ability of endogenously-released serotonin to induce egg laying, we measured the accumulation of unlaid eggs in *ser-1* and *ser-7* null mutant animals. Because *C. elegans* continues to produce eggs even when it cannot lay them, the accumulation of unlaid eggs serves as a convenient measure of defects in egg-laying behavior (Chase and Koelle, 2004). Endogenous serotonin is co-released from the HSN neurons with NLP-3 neuropeptides, and these two signals act semi-redundantly to stimulate egg laying (Brewer et al., 2019). Therefore, the functional role of serotonin in the egg-laying system is best revealed in an *nlp-3* null mutant background: with NLP-3 removed, endogenous serotonin is the strongest remaining signal that stimulates egg laying, and mutations that perturb serotonin signaling thus show much stronger effects on egg laying. This effect is seen in the egg accumulation assays shown in Figure 1E-K and Figure 1-figure supplement 2. Knocking out *tph-1*, the tryptophan hydroxylase enzyme responsible for synthesizing endogenous serotonin (Sze et al., 2000), or knocking out *ser-1* or *ser-7* individually or together, caused only moderate egg-laying defects as seen by accumulation of ∼20-30 unlaid eggs (Figure 1E-H and Figure 1-figure supplement 2A and C). Knocking out *nlp-3* alone, like knocking out serotonin signaling alone, also caused only a modest egg-laying defect in which animals retained ∼23 unlaid eggs (Figure 1E and I, and Figure 1-figure supplement 2A). However, in a *tph-1; nlp-3* double mutant, the worms developed a far more severe egg-laying defect and became bloated with 54.4 ± 2.4 unlaid eggs (Figure 1E and Figure 1-figure supplement 2B).

The above-described results allow us to interpret measurements of animals carrying null mutations for *ser-1* and/or *ser-7* in the *nlp-3* null mutant background. Such animals showed egg-laying defects almost as strong as the defects of the *tph-1; nlp-3* double mutant that completely lacks both serotonin and NLP-3 (Figure 1E, J, and K and Figure 1-figure supplement 2). In the wild type or *nlp-3* null mutant backgrounds, knocking out both SER-1 and SER-7 resulted in a defect not much more severe than knocking out either serotonin receptor alone.

Taken together these data indicate that serotonin signals through the Gα_q_-coupled SER-1 and Gα_s_-coupled SER-7 receptors to initiate egg laying in *C. elegans*. Although these receptors are co-expressed on most muscle cells in the egg-laying system, surprisingly, loss of either the SER-1 or SER-7 receptor resulted in what appeared to be an almost complete loss of the ability of exogenous serotonin to stimulate egg laying and severely disrupted egg laying in response to endogenous serotonin.

### The SER-1 and SER-7 serotonin receptors are each required for endogenous serotonin to coordinate calcium transients in the vm1 and vm2 vulval muscles

We next sought to determine how serotonin signaling through SER-1 and SER-7 induces egg laying in *C. elegans*. We had previously observed that serotonin acts with the neuropeptide NLP-3 to result in simultaneous calcium transients in the vm1 and vm2 vulval muscles. Egg laying only occurs during such simultaneous vm1 and vm2 calcium transients, which drive coordinated contraction of these vulval muscle cells to release eggs (Brewer et al., 2019). We hypothesized that SER-1 and SER-7 are the receptors through which serotonin signals to generate simultaneous vm1 and vm2 calcium transients.

To test this hypothesis, we recorded calcium transients in the vulval muscles of *C. elegans* carrying *ser-1* or *ser-7* null mutations. We expressed the calcium reporter GCaMP5 in the egg-laying muscles and performed one-hour optical recordings of these muscles within freely-behaving animals as previously described (Collins and Koelle, 2013; Brewer et al., 2019).

As controls for the serotonin receptor mutant recordings, we first recorded egg-laying muscle calcium activity in wild-type animals as well as in *tph-1* and/or *nlp-3* null mutant animals. Wild-type animals showed two different types of calcium transients in their vulval muscles: 1) “vm1-only” transients restricted to the vm1 muscles; and 2) “vm1 + vm2” transients that occurred simultaneously in both the vm1 and vm2 muscles (Figure 2A). We never observed a vm2 transient to occur in the absence of a vm1 transient. Wild-type worms had vm1-only transients distributed throughout the entire one-hour recordings (Figure 2B and Figure 2-figure supplement 1). In contrast, vm1 + vm2 transients tended to occur in clusters, known as egg-laying active phases (Waggoner et al., 1998; Brewer et al., 2019), during which a subset of vm1 + vm2 transients were accompanied by release of one or more eggs (Figure 2B and Figure 2-figure supplement 1). In the wild-type, about 17% of the total vulval muscle calcium transients were vm1 + vm2 transients (Figure 3). When *tph-1* (i.e. serotonin) or *nlp-3* were knocked out, there was a modest reduction in the percent of vm1 + vm2 transients (Figure 3B) that correlated with the modest egg laying defects in these mutants (Figure 1E and I, and Figure 1-figure supplement 2A). Knocking out *tph-1* and *nlp-3* together resulted in both an increase in the number of vm1-only transients and a reduction in the number of vm1 + vm2 transients, which combined to produce a significant reduction in the percent of total transients that were of the vm1 + vm2 type (Figure 2B, Figure 2-figure supplement 6, Figure 3). The reduction in the percent of vm1 + vm2 transients correlated with the strong egg-laying defect in the *tph-1; nlp-3* double mutant (Figure 1E and Figure 1-figure supplement 2B). Our recordings in these control genotypes reproduced the findings of Brewer et al. (2019) and confirmed that signaling by serotonin and NLP-3 neuropeptides together lead to the simultaneous activity of the vm1 and vm2 vulval muscles that drives egg laying.

**Figure 2.**
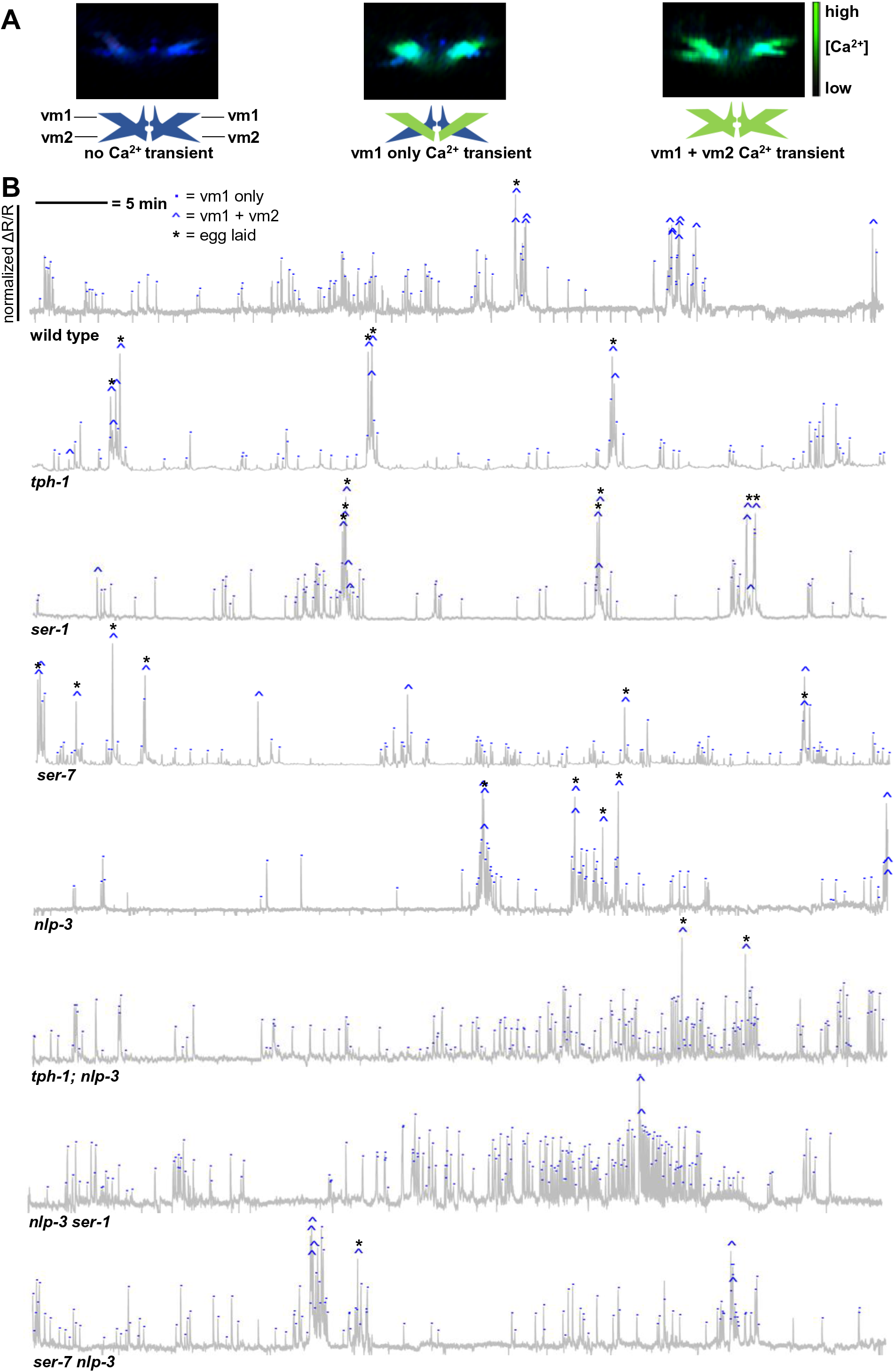
Serotonin signals through SER-1 and SER-7 to coordinate vm1 and vm2 vulval muscle calcium transients. (**A**) Representative still frames from ratiometric calcium imaging of one-hour video recordings of vulval muscles depicting no calcium transient (left), a vm1-only calcium transient (center), and a simultaneous vm1 + vm2 calcium transient (right). The mCherry channel is rendered in blue and the GCaMP channel is superimposed in green, with intensity rendered by ranging from transparent (low calcium) to bright green (high calcium). (**B**) Calcium traces representing one-hour recordings of changes in the GCaMP5/mCherry ratio (ΔR/R) in the vulval muscles of individual worms. Each trace is of a representative animal from the genotype indicated. All traces for each genotype are shown in Figure 2-figure supplements 1-8. Vertical scales have been normalized to the highest peak height within each trace. Each calcium peak was manually scored as vm1-only, marked with a blue dot, or simultaneous vm1 + vm2, marked with a blue caret (^), and transients associated with release of one or more eggs are indicated by asterisks (*).

**Figure 3.**
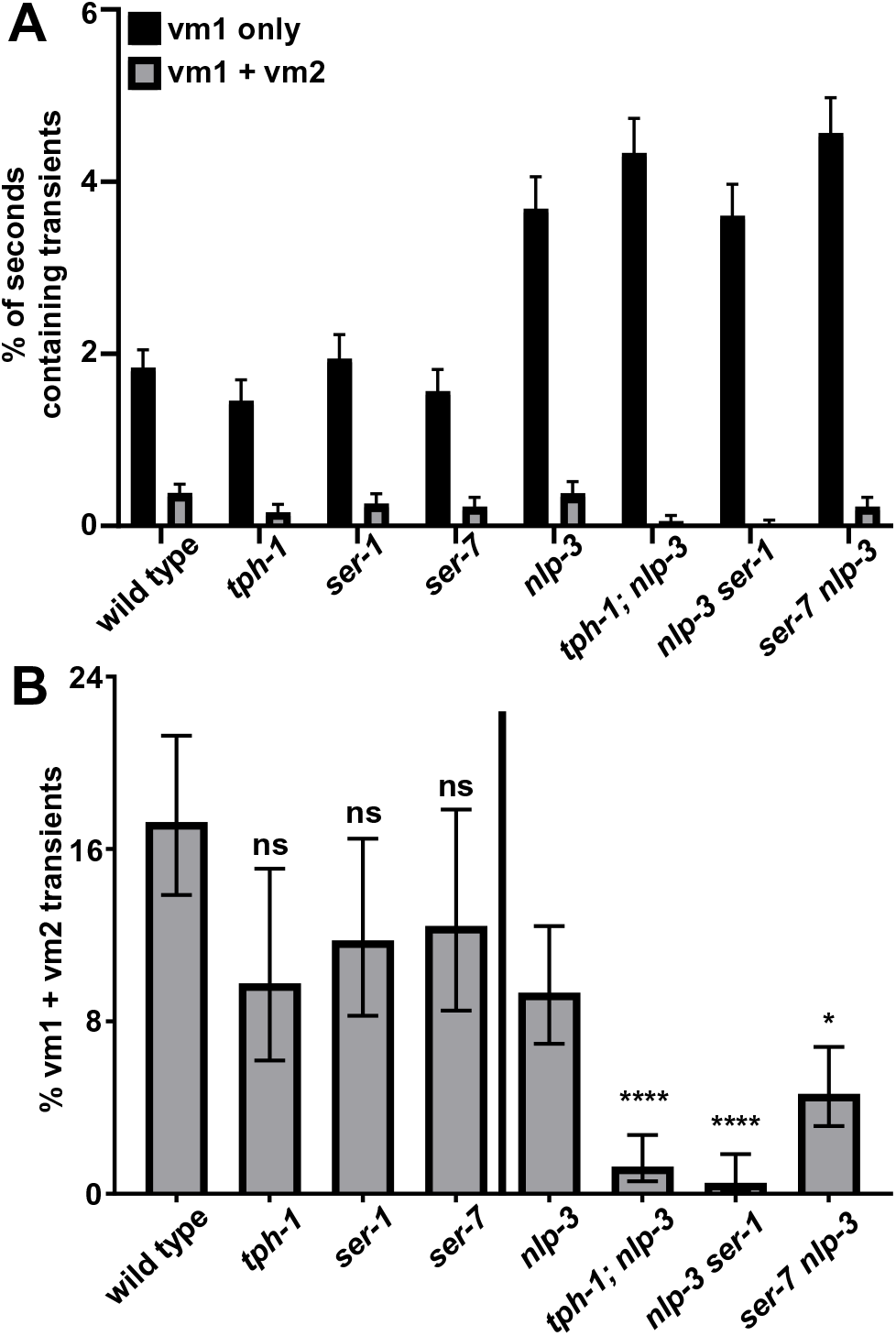
SER-1 and SER-7 are each required for serotonin to signal with NLP-3 to properly induce simultaneous vm1 + vm2 egg-laying muscle contractions. (**A**) Percent of seconds during one-hour recordings that contained either vm1-only or simultaneous vm1 + vm2 calcium transients, averaged from five wild-type recordings and three recordings for each mutant genotype. (**B**) Percentage of total calcium transients that were of the vm1 + vm2 type. Genotypes to the left of the vertical black line were statistically compared to the wild type and genotypes to the right of the vertical black line were statistically compared to *nlp-3*. ns = not significant, * = p<0.05, ** = p<0.01, **** = p<0.0001. Comparing *tph-1;nlp-3* to *nlp-3 ser-1* gave p>0.05 (not significant), whereas comparing *tph-1;nlp-3* to *ser-7 nlp-3* gave p<0.005. All measurements are given with 95% confidence intervals.

Next, we examined the effects of null mutations for SER-1 and SER-7. Single mutants for *ser-1* or *ser-7* each showed a modest reduction in the percentage of vm1 + vm2 transients (Figure 2B, Figure 2-figure supplements 3 and 4, and Figure 3), which is likely responsible for the modest reduction in egg laying seen in these mutants (Figure 1E, G, and H). Crossing the *ser-1* or *ser-7* serotonin receptor null mutants into the *nlp-3* null mutant background isolated serotonin signaling through the remaining serotonin receptor as the remaining driver of egg laying. Both the *nlp-3 ser-1* and *ser-7 nlp-3* double mutants showed a strong reduction in the percentage of vm1 + vm2 transients (Figure 2B, Figure 2-figure supplements 7 and 8, and Figure 3), which correlated with the strong egg-laying defects seen in these double mutants (Figure 1E, J, and K). Indeed, for the *nlp-3 ser-1* double mutant, the defects in egg laying and in the percent of vm1 + vm2 transients were as strong as those of the *tph-1; nlp-3* double mutant (Figure 1E and Figure 3B). The defects in vm1 + vm2 transients in the *ser-7 nlp-3* double mutant were also severe, but slightly less so than those of the *tph-1; nlp-3* double mutant (Figure 3). Together, these data show that endogenous serotonin signals through both the Gα_q_-coupled SER-1 and Gα_s_-coupled SER-7 receptors to coordinate simultaneous vm1 and vm2 vulval muscle transients, and thus egg laying. Knocking out either receptor appears to severely reduce the ability of endogenously-released serotonin to activate the vm2 egg-laying muscles.

### The SER-1 and SER-7 receptors are required on the egg-laying muscles for serotonin to stimulate egg laying

We sought to determine if the SER-1 and SER-7 serotonin receptors are required on the egg-laying muscles themselves to allow serotonin to initiate egg laying. Applying the method developed by Esposito et al. (2007), we used RNAi to cell-specifically knock down genes in the egg-laying muscles. We used the *unc-103e* promoter to drive expression of double-stranded RNA (dsRNA) transcripts specifically in the egg-laying muscles of worms. To ensure that the RNAi effect would remain restricted to the egg-laying muscle cells, these experiments were done in null mutants for SID-1, a double-stranded RNA channel which can allow RNAi to spread from cell to cell (Winston et al., 2002).

We tested the ability of our RNAi system to knock down gene expression specifically in the egg-laying muscles by using animals carrying a transgene that used a 12 kb *ser-7* promoter to express the SER-7 receptor fused to the green fluorescent protein (SER-7::GFP). This transgene expresses SER-7::GFP in all the cells that normally express SER-7, which includes cells in the egg-laying system (Figure 1A; all the egg laying muscle cells and the VC neurons) as well as a set of head neurons (Figure 4A). In these animals, when we used the egg-laying muscle specific promoter to express *ser-7* dsRNA, there was a dramatic loss of SER-7::GFP fluorescence in the egg-laying muscles of 22/25 of worms examined (Figures 4B and D), but no noticeable loss of SER-7::GFP from head neurons (Figure 4B) or VC neurons (Figure 4D), which lie immediately adjacent to the egg-laying muscles. This knockdown was also gene-specific, since expression of a control dsRNA rather than *ser-7* dsRNA did not result in loss of SER-7::GFP fluorescence (Figures 4A and C). We note, however, that our RNAi transgenes may not result in complete knockdown of gene expression, and thus the results described below may reflect partial rather than complete knockdown of gene expression in the egg-laying muscles.

**Figure 4.**
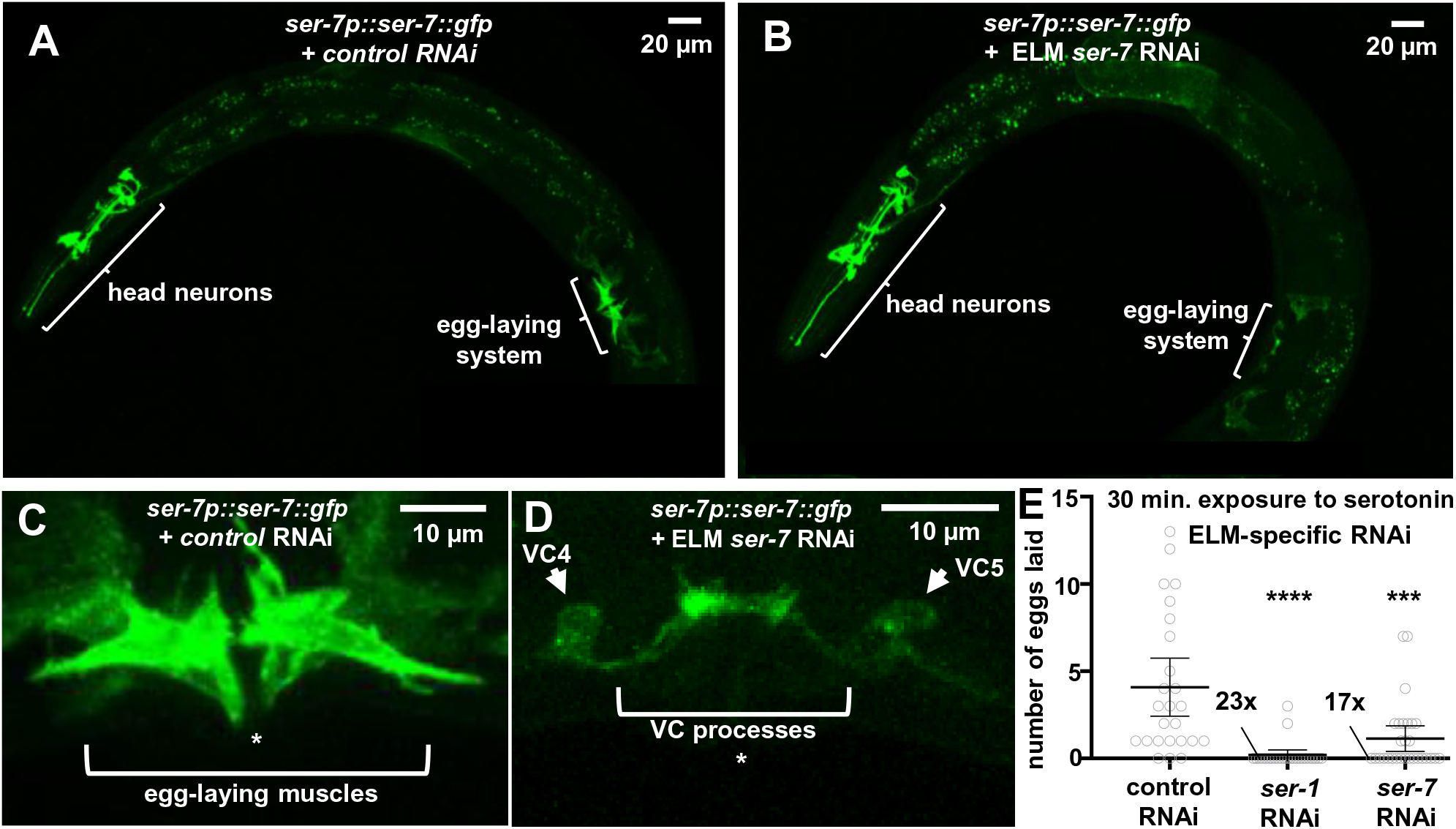
SER-1 and SER-7 are required on the egg-laying muscles for serotonin to stimulate egg laying. (**A**) Image of a transgenic animal in which the *ser-7* promoter (*ser-7p*) drives expression of the SER-7 receptor fused to GFP. GFP fluorescence is seen in a set of head neurons and in the muscles and VC neurons of the egg-laying system. As a negative control, this animal also expresses dsRNA for *ser-1* using an <u>e</u>gg-laying <u>m</u>uscle specific promoter (ELM), which fails to knock down expression of the *ser-7::gfp* transcript, and thus it does not affect GFP fluorescence. (**B**) Image of an animal from the same *ser-7p::ser-7::gfp* strain, but with *ser-7* dsRNA expressed in the egg-laying muscles. Absence of GFP fluorescence in the egg-laying muscles alongside the continued presence of fluorescence in the head neurons indicates successful cell-specific knockdown of *ser-7::gfp*. (**C, D**) Close-up images of the egg-laying system of the same animals shown in (**A**) and (**B**). Knockdown of GFP fluorescence in the egg-laying muscles in (**D**) reveals the fainter GFP fluorescence of the neighboring VC neurons that was obscured in (**C**). Asterisks (*) denote location of the vulval slit. (**E**) Eggs laid per animal after 30 minutes of exposure to exogenous serotonin for the indicated genotypes. RNAi was induced specifically in the egg-laying muscles. n ≥ 25, **** = p<0.0001, *** = p< 0.0005. When many measurements of zero are clustered on the horizontal axis, the number of such data points is indicated. All measurements are given with 95% confidence intervals.

We used the egg-laying muscle-specific RNAi system to knock down either *ser-1* or *ser-7* in the *C. elegans* egg-laying muscles and then tested the ability of exogenous serotonin to induce egg laying. In controls in which neither receptor was knocked down, 22/25 animals laid one or more eggs in response to exogenous serotonin over 30 minutes. In contrast, almost none of the *ser-1* knockdown animals and less than half of the *ser-7* knockdown animals laid any eggs after serotonin treatment, and for both receptor knockdowns the average number of eggs laid was significantly reduced (Figure 4E). To test if *ser-1* and *ser-7* are also required in the egg-laying muscles for endogenous serotonin to stimulate egg laying, we used the same *ser-1* or *ser-7* egg-laying muscle-specific knockdown strains but did not treat with exogenous serotonin and instead simply measured the accumulation of unlaid eggs in adult animals. We saw significant increases in unlaid eggs accumulated for both the *ser-1* and *ser-7* knockdowns (Figure 4-figure supplement 1). The mildness of these increases was expected since whole-animal null mutants for these receptors have relatively mild effects (Figure 1E). These cell-specific RNAi results show that both SER-1 and SER-7 are required on the egg-laying muscles for serotonin to properly induce egg laying.

### Gα_q_ and Gα_s_ signaling are required in the egg-laying muscles to combine endogenous signals that stimulate egg laying

We next investigated whether the G proteins through which SER-1 and SER-7 signal, Gα_q_ and Gα_s_, respectively, are necessary in the egg-laying muscles for proper egg laying in response to endogenous signals within the animal. We used our RNAi system to knock down the genes for Gα_q_ and Gα_s_ specifically in the egg-laying muscles and measured the accumulation of unlaid eggs.

We found that RNAi knockdown of Gα_q_ or Gα_s_ in the egg-laying muscles had mild but significant effects on the accumulation of unlaid eggs, but that knocking down both Gα proteins together had a stronger effect (Figure 5). This strong defect was not the result of the Gα_q_/Gα_s_ double knockdown causing developmental defects in the egg-laying muscles, since a) the egg-laying muscle-specific *unc-103e* promoter used to express dsRNA for these gene knockdowns only turns on in the egg-laying muscles as the muscle cells are completing their terminal differentiation (Ravi et al., 2018); and b) we labeled the egg-laying muscles with a fluorescent protein in Gα_q_/Gα_s_ double knockdown animals and saw no visible morphological defects in these muscle cells (Figure 5-figure supplement 1). The mild defects seen in the single Gα knockdowns are difficult to interpret, since we are not certain of the extent to which RNAi reduced the levels of the Gα proteins. Even so, we note that the strong egg-laying defect that resulted from Gα_q_/Gα_s_ double knockdown in the egg-laying muscles (Figures 5) is stronger than the egg-laying defects observed in animals with complete knockouts of both *ser-1* and *ser-7,* or in animals with a *tph-1* knockout that completely eliminates endogenous serotonin (Figure 1E, G, and H and Figure 1-figure supplement 2A and 2C). Therefore, serotonin appears not to be the only signal that generates Gα_q_ and Gα_s_ activity in the egg-laying muscles to stimulate egg laying. These data indicate that there must be other GPCRs on the egg-laying muscles, in addition to SER-1 and SER-7, that signal through Gα_q_ and Gα_s_ to stimulate egg laying. Therefore, normal levels of egg-laying activity result from Gα_q_ and Gα_s_ acting in the egg-laying muscles to combine signals from SER-1, SER-7, and additional GPCRs.

**Figure 5.**
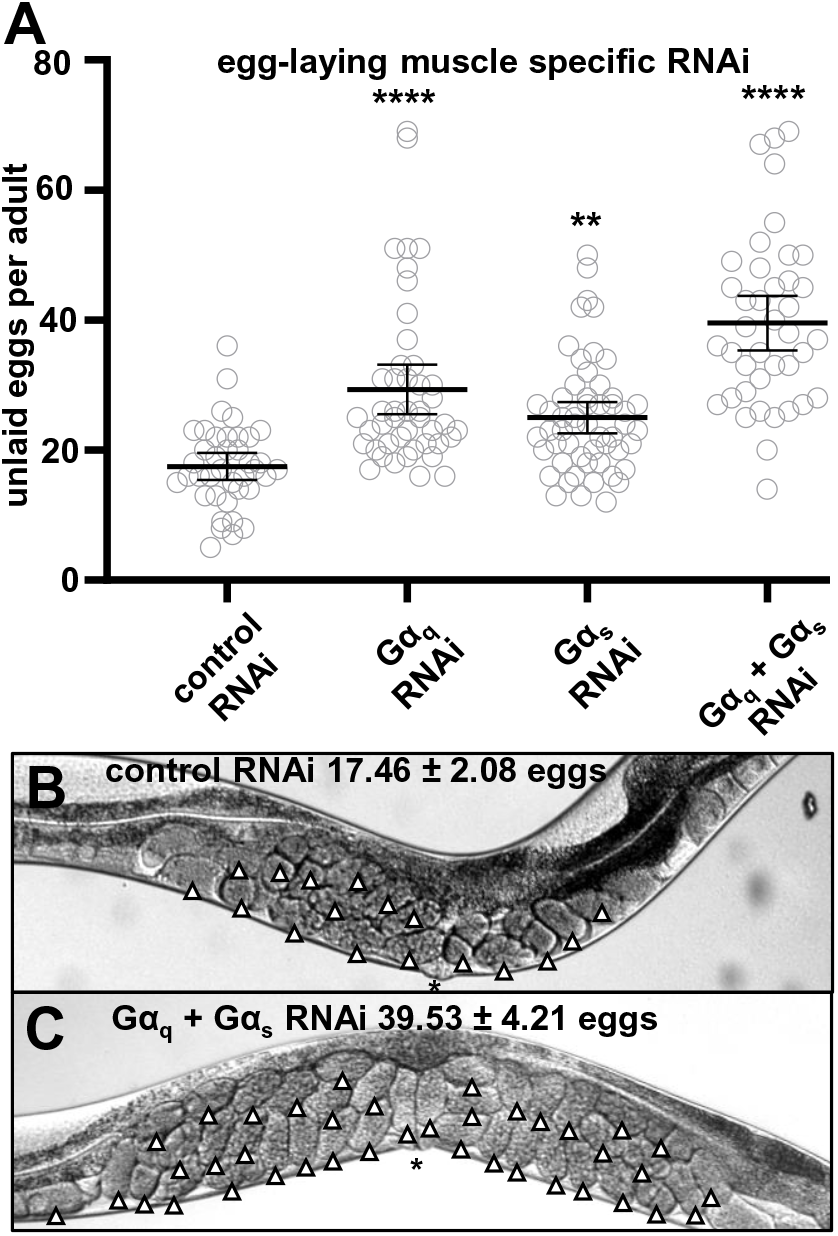
Gα_q_ and Gα_s_ signaling are necessary in the egg-laying muscles to drive proper egg laying. Egg-laying muscle specific RNAi was used to knock down Gα proteins and the resulting accumulation of unlaid eggs was measured. Anti-*gfp* RNAi was used as a negative control. (**A**) Average number of unlaid eggs per adult worm, n≥30 for each genotype. ** indicates p<0.01, **** indicates p<0.0001. (**B, C**) Photographs of worms of the indicated genotypes, with unlaid eggs indicated by white arrowheads. The vulval slit is indicated by *. All measurements are given with 95% confidence intervals.

### Overexpressed SER-1 is sufficient to allow serotonin to induce egg laying in the absence of SER-7

Results presented above show that knocking out or knocking down either the SER-1 or SER-7 serotonin receptors result in severe defects in the ability of serotonin to induce egg laying. In some cases, the defects observed were as strong as those caused by knocking out both SER-1 and SER-7 at the same time or as strong as those seen when completely eliminating serotonin with a *tph-1* null mutation (Figure 1E). These results raise the question of whether serotonin absolutely requires both SER-1/Gα_q_ and SER-7/Gα_s_ signaling to induce egg laying, or whether these two signaling pathways might rather combine to induce egg laying in a more nuanced fashion. Thus, we designed several different experiments to determine if increasing the strength of just one of the two pathways could induce egg laying in the absence of the other pathway.

The first method was to simply overexpress one serotonin GPCR by increasing the copy number of the GPCR gene. Previous genetic studies have shown that overexpression can increase the normal functions of a GPCR in a manner that is suppressed by knocking out the endogenous ligand for that GPCR (Ringstad and Horvtiz, 2008; Harris et al., 2010; Brewer et al., 2019; Fernandez et al., 2020), suggesting that the overexpressed GPCR is activated by its endogenous ligand to signal at a higher level than would the endogenous levels of the GPCR. Indeed, overexpressing SER-1 in *C. elegans* was shown to increase egg laying in a manner completely dependent on endogenous serotonin (Fernandez et al., 2020).

To overexpress serotonin receptors, we used chromosomally-integrated transgenes that carry multiple copies of the complete *ser-1* or *ser-7* genes, including their own promoters, resulting in overexpression of these genes in the same cells that normally express them (Fernandez et al., 2020). We tested the ability of exogenous serotonin to induce egg laying in animals overexpressing one serotonin receptor while also carrying a deletion mutation for the other serotonin receptor. Our results are graphed in Figure 6, and the design and logic of this experiment are schematized in Figure 6-figure supplement 1. We found that animals overexpressing *ser-7* in a *ser-1*-null background were not able to lay eggs in response to exogenous serotonin, similar to animals that simply lacked *ser-1.* However, animals overexpressing *ser-1* in a *ser-7*-null background did lay eggs in response to exogenous serotonin, unlike animals that simply lacked *ser-7*.

**Figure 6.**
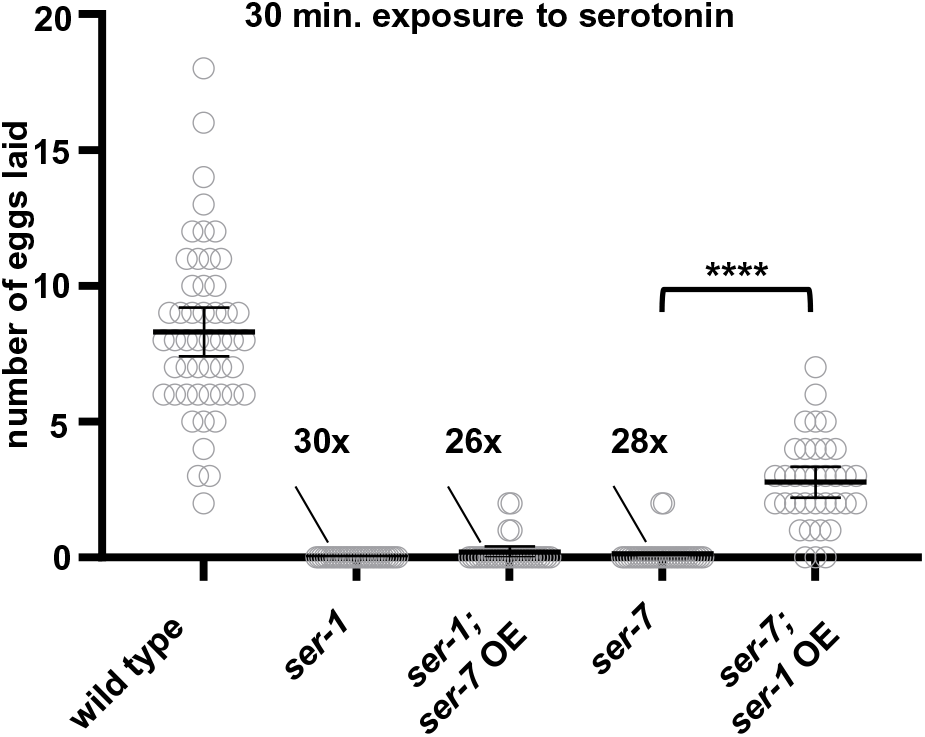
Over-expression of SER-1 is sufficient to allow serotonin to induce egg laying in the absence of SER-7. Number of eggs laid per worm during a 30-minute exposure to exogenous serotonin on NGM plates. “OE” indicates overexpression of the indicated receptor from a high-copy transgene that carries the entire receptor gene, including its promoter, so that the receptor is overexpressed in the same cells that normally express the endogenous receptor. For three genotypes, the number of animals that laid zero eggs are indicated since the many individual data points bunched on the horizontal axis are difficult to discern. n ≥30, **** = p<0.0001. All measurements are given with 95% confidence intervals.

These results show that while SER-1/Gα_q_ signaling is normally not sufficient to allow serotonin to induce egg laying in the absence of SER-7/Gα_s_ signaling, artificially increasing SER-1 expression levels overcomes this limitation. It is difficult to interpret the negative result from the converse experiment, in which overexpressed SER-7 failed to induce egg laying in the absence of SER-1. It could be that our SER-7 overexpression experiment may have not increased SER-7/Gα_s_ signaling to a high enough level to induce egg laying in the absence of SER-1/Gα_s_ signaling or it could mean that SER-7 alone is incapable of driving egg-laying. Thus, we devised additional methods, described below, to artificially increase Gα_q_ and/or Gα_s_ signaling in the egg-laying muscles and determine which pathway(s) are sufficient to drive egg laying.

### Gα_q_ or Gα_s_ signaling in the egg-laying muscles is sufficient to drive egg laying

A previous study developed a DREADD (<u>D</u>esigner <u>R</u>eceptor <u>E</u>xclusively <u>A</u>ctivated by <u>D</u>esigner <u>D</u>rugs; Lee et al., 2014) to activate Gα_q_-signaling in *C. elegans* in response to the drug clozapine N-oxide (CNO) (Prömel et al., 2016). We acutely activated Gα_q_-signaling specifically in the egg-laying muscles by transgenically expressing this designer Gα_q_-coupled receptor using the egg-laying muscle specific promoter and treating the worms with CNO. This induced egg laying (Figure 7A and B). In contrast, worms carrying a control transgene were unable to lay eggs in response to CNO (Figure 7A).

**Figure 7.**
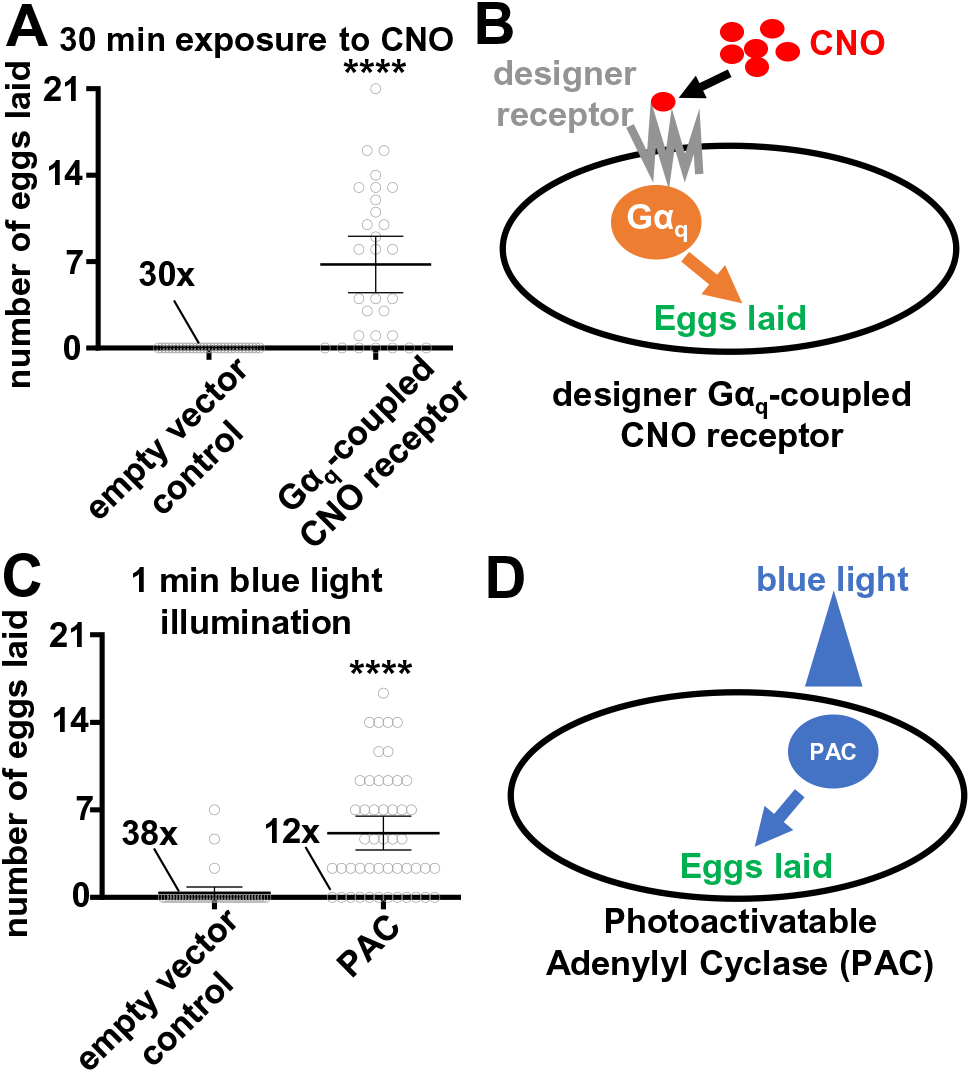
Signals from either the Gα_q_ or Gα_s_ pathways in the egg-laying muscles can be sufficient to drive egg laying. (**A**) Number of eggs laid per worm after 30 minutes of exposure to 10 mM CNO in worms expressing either the designer Gα_q_-coupled CNO-responsive receptor in their egg-laying muscles or carrying a control empty vector transgene. (**B**) Schematic of the design of the experiment shown in (**A**). (**C**) Number of eggs laid per worm after 1 minute of exposure to blue light in animals expressing <u>P</u>hotoactivatable <u>A</u>denylyl <u>C</u>yclase (PAC) in the egg-laying muscles to induce the downstream effects of Gα_s_ signaling. (**D**) Schematic of the design of the experiment shown in (**C**). For both assays n ≥30. **** = p<0.0001. All measurements are given with 95% confidence intervals.

Next, we determined if Gα_s_-signaling in the egg-laying muscles was sufficient to drive egg laying. To date there is no designer Gα_s_-coupled receptor that is functional in *C. elegans* (Prömel et al., 2016) and we were unsuccessful in further attempts to design such a receptor (data not shown). Gα_s_ signals by activating adenylyl cyclase, which in turn generates cAMP. A photoactivatable adenylyl cyclase (PAC) has been successfully used in the cholinergic neurons and body wall muscles of *C. elegans* to evoke changes in locomotion (Steuer Costa et al., 2017; Henss et al., 2022). We generated transgenic animals that express PAC in their egg-laying muscles and found that blue light activation of PAC was able to induce egg laying in these worms, whereas control worms carrying an empty vector transgene were unable to lay eggs in response to blue light (Figure 7C and D, and Video 1). These results demonstrate that activation of either the Gα_q_ or Gα_s_ pathways in the egg-laying muscles is sufficient to induce egg laying, and that these G protein signals can originate from sources other than a serotonin receptor.

### The combination of two subthreshold signals from different Gα_q_-coupled receptors in the egg-laying muscles is sufficient to drive egg laying

The results above demonstrate that artificially-induced Gα_q_ or Gα_s_ signaling in the egg-laying muscles can be sufficient to induce egg laying. However, we also found that neither endogenous SER-1/Gα_q_ signaling alone nor endogenous SER-7/Gα_s_ signaling alone in these same egg-laying muscles is sufficient to drive egg laying; instead, both these endogenous signaling pathways must be active at the same time to induce egg laying. To reconcile these findings, we hypothesized that the endogenous levels of SER-1/Gα_q_ and SER-7/Gα_s_ signaling are both “subthreshold,” i.e., occur at low levels that are not sufficient to properly activate egg laying on their own, and together sum to reach the threshold necessary to activate egg laying.

To test this hypothesis, we generated an artificial subthreshold G protein signal that was unable to activate egg laying on its own and determined if it was capable of activating egg laying when combined with another subthreshold G protein signal. The designer Gα_q_-coupled receptor offered the potential to tune the levels of Gα_q_ signaling it induces: we titrated the concentration of its CNO ligand to find a concentration (2 mM) that was just below the threshold required to activate egg laying on its own (Figure 8A). We then expressed the designer Gα_q_-coupled receptor in the egg-laying muscles of worms lacking the SER-7 receptor, so that serotonin could only signal to induce egg laying via SER-1/Gα_q_ (Figure 8C). Exposure to either exogenous serotonin or to 2 mM CNO was unable to induce egg laying, as expected. However, when both exogenous serotonin and 2 mM CNO were applied to the worm at the same time, these two subthreshold Gα_q_-coupled signals combined to activate egg laying (Figures 8B and C).

**Figure 8.**
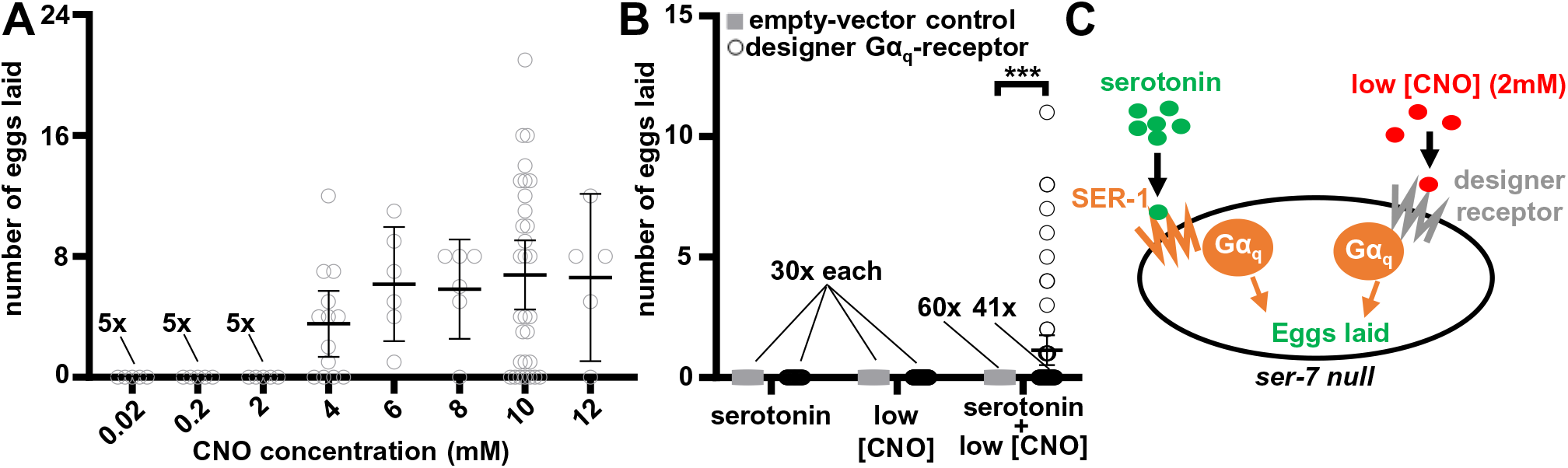
The combination of subthreshold signals from two different Gα_q_-coupled receptors in the egg-laying muscles is sufficient to drive egg laying. (**A**) Number of eggs laid after 30 minutes by worms expressing the designer Gα_q_-coupled CNO receptor in their egg-laying muscles after being treated with a range of CNO concentrations. n≥5. (**B**) *ser-7* mutant animals, with or without expression of the CNO receptor in their egg-laying muscles, were exposed to either 25 mM exogenous serotonin, 2 mM CNO, or both. Eggs laid after 30 minutes were counted. n≥30. In each condition, the number of animals that laid zero eggs is indicated above the data point. *** = p<0.001. **C**) Schematic of the design and results of the experiment shown in (B). Note that the lack of the SER-7 receptor results in serotonin signaling only via SER-1/Gα_q_. The threshold of signaling required to activate egg laying is reached by combining two subthreshold signals generated by: 1) partially activating the designer Gα_q_-coupled receptor with a low concentration of 2mM CNO, and 2) activating the Gα_q_-coupled SER-1 receptor with exogenous serotonin. All measurements are given with 95% confidence intervals.

## Discussion

The principal finding of this study is that the SER-1 and SER-7 serotonin receptors, as well as additional Gα_q_ and/or Gα_s_ coupled receptors, all signal together in the *C. elegans* egg-laying muscles to help induce their coordinated contraction and thus the laying of eggs. While signaling from endogenous levels of just one of these receptors alone is not strong enough to induce egg laying, together the signals from multiple types of GPCRs on the same cells combine to reach a threshold that can activate egg laying (Figure 9). This study is perhaps the most detailed to date of how cells within an intact organism integrate signaling by multiple GPCRs to generate a concerted response to the complex mixture of chemical signals impinging upon them. Such signal integration is a challenge faced by virtually all cells within multicellular organisms, and the findings from our study of how this is accomplished in the *C. elegans* egg-laying muscles likely generalize to similar situations faced by other cells.

**Figure 9.**
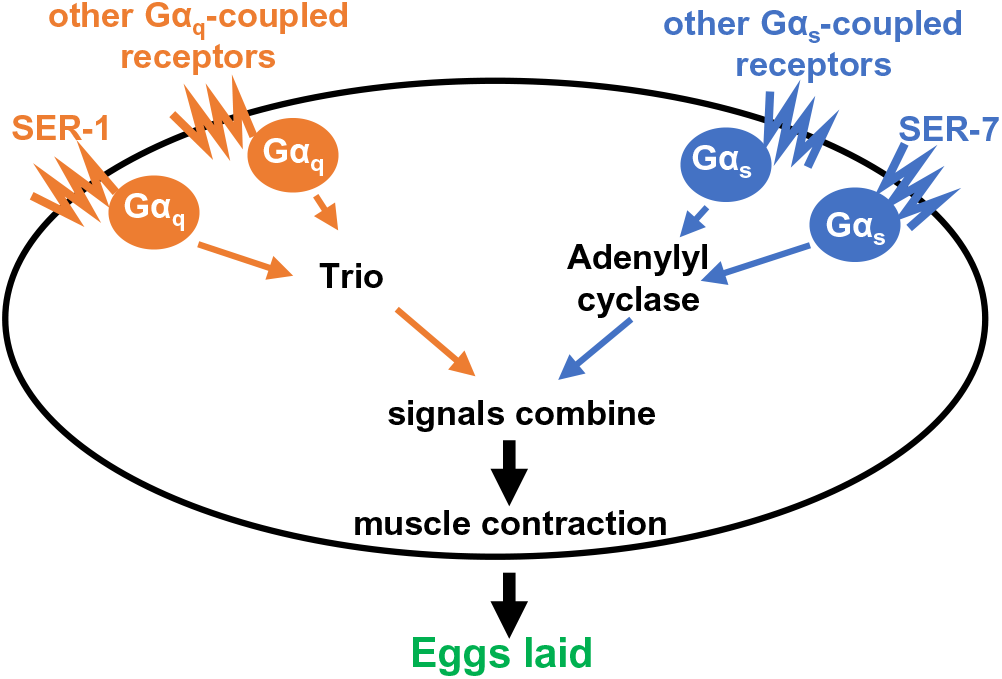
Model for the regulation of egg-laying muscle activity by multiple Gα_q_- and Gα_s_-coupled GPCRs. Each Gα protein is activated in parallel by a serotonin receptor and additional GPCRs. Gα_q_ directly activates the RhoGEF protein Trio, while Gα_s_ directly activates adenylyl cyclase. The two signaling pathways intersect at a downstream point, yet to be determined, to promote muscle contraction, which drives egg laying. Individual GPCRs provide signaling too weak to result in a significant increase in egg laying, but combining signaling by multiple GPCRs through Gα_q_, Gα_s_, or both can result in a significant behavioral outcome.

### Multiple GPCRs signal through Gα_q_ and Gα_s_ to activate excitable cells

We found that knocking down both Gα_q_ and Gα_s_ in the egg-laying muscles resulted in a dramatic defect in egg laying, while loss of the SER-1 and SER-7 serotonin receptors that activate these G proteins, either from the entire animal or from the egg-laying muscles alone, only had a modest effect. Therefore, serotonin appears to combine with other endogenous signals to generate sufficient Gα_q_ and Gα_s_ signaling in the egg-laying muscles to induce egg laying. Treating animals with a high concentration of exogenous serotonin is sufficient to induce egg laying, and, even in this artificial situation, both the SER-1 and SER-7 receptors must operate in parallel on the egg-laying muscles to mediate this effect, as loss of either receptor from the egg-laying muscles results in almost complete loss of the ability of exogenous serotonin to induce egg laying. We were able to generate artificial circumstances in which activation of a single type of GPCR on the egg-laying muscles (either overexpressed SER-1 or the designer CNO receptor) could induce egg laying. Additionally, activation of the signaling pathway downstream of SER-7 with a photoactivatable adenylyl cyclase was sufficient to induce egg laying. Nonetheless, our results show that the normal situation in wild-type animals is that egg laying is induced by the combined signaling from multiple Gα_q_-and Gα_s_-coupled receptors.

What other signals besides serotonin might be acting on the egg-laying muscles to promote egg laying? The neuropeptide NLP-3 is co-released with serotonin onto the egg-laying muscles to promote egg laying (Brewer et al., 2019), and it is possible that the NLP-3 receptor, which has not yet been identified, is one of the additional GPCRs expressed on these muscles. If so, the NLP-3 receptor would combine its effects with those of the SER-1 and SER-7 serotonin receptors to induce egg laying. NLP-3 signaling on its own, like signaling from the serotonin receptors, produces modest effects, with dramatic defects in egg laying seen only when both NLP-3 and serotonin signaling are lost simultaneously. A systematic analysis of the expression of all *C. elegans* neurotransmitter GPCRs on the egg-laying muscles (Fernandez et al., 2020) found that, besides SER-1 and SER-7, three additional Gα_q_- and Gα_s_-coupled GPCRs are expressed on these cells: the dopamine receptor DOP-4, the tyramine receptor TYRA-3, and the metabotropic acetylcholine receptor GAR-3. Just as for SER-1 and SER-7, knockouts for any one of these receptors have, at most, modest effects on the accumulation of unlaid eggs (Fernandez et al., 2020), consistent with the hypothesis that the G proteins Gα_q_ and Gα_s_ integrate signals from a variety of GPCRs on the egg-laying muscles to maintain proper egg laying.

Our finding that multiple GPCRs combine signaling in the egg-laying muscles of *C. elegans* to induce strong behavioral effects appears to be a general feature of GPCR signaling in excitable cells within multicellular organisms. Heterotrimeric G protein signaling in *C. elegans* neurons and muscles has been studied intensively for decades (reviewed by Koelle, 2018). These studies have included a variety of forward genetic screens for mutants with various behavioral defects resulting from disruption of G protein signaling (Trent et al., 1983; Desai and Horvitz, 1989; Bargmann et al., 1993; Miller et al., 1996; Bany et al., 2003). Together, these screens have been carried out on a saturation scale such that mutations in perhaps every gene involved in heterotrimeric G protein signaling have been sampled to identify those that are critical to control the behaviors studied. These screens have produced a large set of mutants for the G proteins themselves, the Regulators of G protein Signaling (RGS proteins) that terminate signaling, and the proteins that act downstream of the G proteins. However, mutants for GPCRs are almost absent from the results of these screens (Koelle, 2018). There is also a conspicuous paucity of GPCR mutations that have arisen from the century of forward genetic screens that have been carried out in *Drosophila*, despite GPCRs constituting the single largest family of proteins in metazoan organisms such as worms and flies (Hanlon and Andrew, 2015). One possible explanation for this paradox could be that GPCR mutations are generally lethal, however, GPCR knockout mutations generated in these model invertebrates are rarely, if ever, lethal and typically do not show overtly obvious behavioral defects (e.g. Fernandez et al., 2020). Thus, the near absence of GPCR mutants arising in *C. elegans* or *Drosophila* behavioral genetic screens suggests that loss of a single neurotransmitter or neuropeptide GPCR rarely causes significant defects in the behaviors that have been studied, even though mutations in heterotrimeric G proteins show severe defects in the control of these same behaviors. This paradox can be resolved if, as in egg laying, multiple GPCRs combine their signaling to result in behavioral effects.

Studies of G protein signaling in vertebrate cardiomyocytes (heart muscle cells) parallel our finding in *C. elegans* egg-laying muscles that multiple co-expressed GPCRs together regulate muscle contractility. A number of neurotransmitters and neuropeptides modulate heart muscle function, and their roles in heart disease have prompted studies of the GPCRs on cardiomyocytes that mediate the effects of these signals (Wang et al., 2018; Lymperopoulos et al., 2021). These include four receptors that mediate signaling by epinephrine and norepinephrine: the Gα_s_-coupled β1- and β2-adrenergic receptors (Bristow et al., 1986), and the Gα_q_-coupled α1a- and α1b-adrenergic receptors (McCloskey et al., 2003; O’Connell et al., 2003). Cardiomyocytes also express Gα_q_-coupled receptors for the peptide hormones vasopressin (Xu and Gopalakrishnan, 1991) and angiotensin II (Meggs et al., 1993). While it is clear that together these signals and receptors regulate heart muscle contractility and that each plays crucial roles mediating heart disease, it is less clear how these multiple signals and receptors combine their effects within the intact organism to orchestrate proper control of heart muscle function. The *C. elegans* egg-laying muscles provide a more easily manipulatable genetic model system for understanding how multiple GPCR signals together regulate muscle function.

### How do Gα_q_ and Gα_s_ signals combine to modulate activity of excitable cells?

In the *C. elegans* egg laying circuit, our results suggest serotonin released by the HSNs acts directly on the egg-laying muscles to make these muscle cells more excitable, enabling other signals that depolarize the muscle cells to trigger the simultaneous vm1 + vm2 muscle cell contractions that release eggs. Previous work has identified other signals released by cells other than the HSNs onto the egg-laying muscles to depolarize the muscle cells and act as the final trigger for contraction. The VC neurons release acetylcholine at synapses onto the vm2 egg-laying muscle cells (see Figure 1), which acts via acetylcholine-gated ion channels (i.e. nicotinic receptors) to excite the vm2 cells (Kopchock et al., 2021). The vm1 muscle cells separately receive an as-yet uncharacterized excitatory signal during every body bend (Collins and Koelle, 2013), but this signal does not trigger egg-laying muscle contractions unless the HSN neurons have first released serotonin and/or NLP-3 neuropeptides (Brewer et al., 2019). Our results in this work show that serotonin enables contraction responses in the egg-laying muscles by acting via both the Gα_q_-coupled SER-1 and Gα_s_-coupled SER-7 receptors.

Studies in vertebrate heart muscle suggest mechanisms by which Gα_q_ and Gα_s_ signaling may together promote muscle contraction. Gα_s_ signaling is proposed to promote contraction through a complex mechanism in which Gα_s_ activates adenylyl cyclase to produce cAMP, which in turn binds and activates protein kinase A (PKA), causing PKA to phosphorylate a variety of targets that promote contractility (reviewed by Salazar et al., 2007). The proposed effects of PKA include phosphorylating the L-type Ca^2+^ channel to promote extracellular Ca^2+^ entry into the muscle cell, phosphorylating the ryanodine receptor (a Ca^2+^ channel on internal membranes) to promote Ca^2+^ release from internal stores, and phosphorylating the Ca^2+^-binding muscle filament protein troponin to promote the ability of Ca^2+^ to trigger contraction. Gα_q_ signaling has complex effects on vertebrate heart function, including some that could combine with Gα_s_ signaling to promote muscle contraction (Lin et al., 2001; McCloskey et al., 2003). First, Gα_q_ activates its effector phospholipase C to ultimately lead to phosphorylation of Gα_s_-coupled β-adrenergic receptors, altering their mode of signaling and thus the ability of epinephrine to regulate muscle contraction (Wang et al., 2018). Second, Gα_q_ and Gα_s_ signaling can collaborate to activate IP_3_ receptors which, like ryanodine receptors, are Ca^2+^ channels that release Ca^2+^ from internal stores to promote muscle contraction. In this mechanism, Gα_q_ directly activates the enzyme phospholipase C (Smrcka et al., 1991; Taylor et al., 1991), which generates the second messenger IP_3_ that directly binds and activates the IP_3_ receptor. Gα_s_ signaling, as noted above, activates the protein kinase PKA, which can phosphorylate and activate IP_3_ receptors (Taylor, 2017).

The mechanism by which Gα_q_ and Gα_s_ signaling alter muscle and neuron function has been independently addressed through other studies in *C. elegans*. In the egg-laying muscles, genetic studies show Gα_q_ promotes contraction mainly not via phospholipase C (Dhakal et al., 2022), as suggested by vertebrate heart muscle studies (Salazar et al., 2007), but rather by activating the other major Gα_q_ effector, the RhoGEF protein Trio, which in turn activates the small G protein Rho (Chikumi et al., 2002; Lutz et al., 2005; Lutz et al., 2007, Rojas et al., 2007; Williams et al., 2007). The different conclusions reached in vertebrate heart versus *C. elegans* egg-laying muscles may reflect differences in how Gα_q_ regulates these two types of muscles or rather could reflect the different experimental approaches used to study these two systems.

Our work, as well as previous studies in *C. elegans*, have also examined the relationship between Gα_q_ and Gα_s_ signaling. Studies of neurons in the *C. elegans* locomotion circuit suggest that Gα_q_ signaling provides a core function that activates neuron output while Gα_s_ signaling potentiates a downstream step in Gα_q_ signaling. For example, hyperactivation of the Gα_q_ pathway could rescue locomotion defects seen in Gα_s_ reduction of function mutants, yet hyperactivation of the Gα_s_ pathway could not rescue locomotion defects seen in Gα_q_ reduction of function mutants (Reynolds et al., 2005). Our work shows that activating high enough levels of either Gα_q_ or Gα_s_ signaling alone in the *C. elegans* egg-laying muscles is sufficient to promote egg laying. However, our studies were carried out within intact animals in which low levels of both Gα_q_ and Gα_s_ signaling may occur in the background as we hyperactivate one of these signaling pathways. We also used RNAi to knock down either Gα_q_ or Gα_s_ in the egg laying muscles and observed only partial inhibition of egg laying, but we cannot ensure that knockdown of the Gα proteins was complete. Because null mutations in Gα_q_ and Gα_s_ are lethal in *C. elegans* (Reynolds et al., 2005), it is not straightforward to generate more rigorous genetic studies of the relationship of these two signaling pathways.

## Conclusions

Combining signaling by multiple G protein coupled receptors appears to be a universal mechanism used to modulate activity of neurons and muscle cells in multicellular organisms. The logic of why multiple GPCRs are found on a single cell and how these GPCR signals can meaningfully funnel through just a few types of Gα proteins has long been a mystery. Our work shows that within an intact animal, multiple Gα_q_- and Gα_s_-coupled receptors co-expressed on the same cells each generate weak signals that individually have little effect but sum together to produce enough signaling output to impact behavior. This system allows a cell to gather multiple independent pieces of information from the complex soup of chemical signals in its environment and compute an appropriate response. In the case of the *C. elegans* egg-laying system, the multiple neurotransmitters and neuropeptides released by the egg-laying circuit are sensed to determine when conditions are right for the animal to lay an egg. More generally, this system for computing outcomes by integrating multiple inputs provides neurons and muscles with a vastly flexible mechanism for processing information.

## Materials and Methods

### Strains and culture

A complete list of the *C. elegans* strains and transgenes used in this paper is found in Supplementary File 1. *C. elegans* were maintained at 20°C on standard nematode growth media (NGM) seeded with OP50 strain of *Escherichia coli* as their food source. Mutants and animals carrying chromosomally-integrated transgenes were backcrossed 2-10x to N2 (wild type) to generate clean genetic backgrounds, as indicated in Supplementary File 1. New strains were constructed using standard genetic cross procedures and genotypes were confirmed by PCR genotyping or sequencing. Extrachromosomal array transgenic strains were generated through microinjection. Phenotypes were typically scored in animals from ≥5 independent transgenic lines, and at least one independent line has been frozen for storage. Supplementary File 1 details the strains used and how each transgenic strain was constructed.

### Molecular biology

The construction of the plasmids used in this manuscript are described in Supplementary File 1.

### Egg-laying muscle specific RNAi

Transgenic animals with egg-laying muscle specific RNAi were created as described in Esposito et al. (2007). PCR was used to fuse the *unc-103e* promoter upstream of an exon-rich region of the gene target by RNAi. To increase the yield of the fusion PCR product, NEBuilder HIFI DNA Assembly Mix (NEB) was used to fuse the promotor fragment to the exon-rich gene fragment prior to nested PCR. Two fusion PCR products for each gene of interest were injected into C. elegans, one expressing sense RNA and the other antisense RNA. The sense and antisense RNA strands anneal in the cell to form the dsRNA used for RNAi. Due to the highly identical sequences of Gα_q_ and Gα_s_, care was taken to choose dissimilar regions of the genes encoding Gα_q_ and Gα_s_ to target with RNAi. The regions chosen had no more than 14 bp of contiguous sequence identity. 50-100 ng/µl of fusion PCR product expressing sense RNA and 50-100 ng/µl of fusion PCR product expressing antisense RNA were injected into *sid-1(qt9) V; lin-15(n765ts) X* animals along with 10ng/ul pCFJ90 (pharyngeal mCherry co-injection marker), 50ng/ul pL15EK (*lin-15(+)* co-injection marker), and 25ng/ul DH5alpha genomic DNA digested with BamHI/HindIII. The *sid-1(qt9)* mutation kept the RNAi cell-specific by preventing cell-to-cell spreading of the RNAi via systemic RNAi. Supplementary File 1 details the construction of the fusion PCR products, including the exact concentrations injected for each DNA and the sequences used for the *unc-103e* promoter region and each exon-rich gene region that was targeted by RNAi. During the knockdown of G proteins, mCherry was expressed in the egg-laying muscles to demonstrate that the G protein knockdown did not interfere with muscle development (Figure 5–supplement 1). There was no statistical difference between the number of eggs retained in animals in expression (data not shown).

### Calcium imaging

Animals were staged as late-L4 larvae and recorded 24 hours later. Freely-behaving animals were mounted between a glass coverslip and a ∼1 cm^2^ chunk from an NGM plate containing OP50 food for imaging as previously described (Collins and Koelle, 2013, Collins et al., 2016, Ravi et al., 2018). A brightfield and two fluorescence channels (for the green GCaMP calcium sensor and a control mCherry protein) were recorded with a 20X air objective using a Zeiss LSM 880 microscope. Recordings were collected at ∼16 fps at 256 x 256 pixel, 16 bit resolution, for 1 hour. Three 1-hour recordings were collected for each genotype studied. As previously described (Brewer et al., 2019), calcium imaging was recorded in both the vm1 and vm2 vulval muscles simultaneously and ratiometric analysis of the calcium recordings was performed in Volocity (PerkinElmer) to generate traces of calcium transients. As described in Brewer et al. (2019), a video of each peak was examined and scored as vm1-only or vm1 + vm2.

### Confocal imaging

Animals were mounted on microscope slides with 2% agarose pads containing 120 mm Optiprep (Sigma Millipore) to reduce refractive index mismatch (Boothe et al., 2017) and a 22×22–1 microscope cover glass (Fisher Scientific) was placed on top of the agarose pad. Animals were anesthetized using a drop of 150 mM sodium azide (Sigma Millipore) with 120 mm Optiprep. Z-stack confocal images of *C. elegans* staged 24 hours post L4 were taken on a Zeiss LSM 880 microscope using a 40X water-immersion objective lens.

### Serotonin-induced egg laying on NGM plates

This assay was adapted from the work of Hobson et al. (2006). NGM plates containing 26 mM serotonin creatine sulfate monohydrate (Sigma, H7752-5G) were poured and seeded with OP50 one day prior to assay. Animals were staged as late L4 larvae for assay 24 hours later. At time 0 of the assay, 5-10 worms were placed on the serotonin plates, spaced in a manner that it was unlikely they would be able to crawl near each other prior to being paralyzed by the serotonin. Serotonin induced paralysis, which resulted in the worms remaining adjacent to the eggs they laid during the time course, making it was possible to attribute the number of eggs laid to each individual worm.

### Serotonin- or CNO-induced egg laying in M9 buffer

Animals were staged as late L4 larvae for assay 24 hours later. Serotonin creatine sulfate monohydrate from Sigma (H7752-5G) and Clozapine N-oxide dihydrochloride (CNO) from Fisher Scientific (sourced from Tocris Bioscience (6329/10)) were dissolved to desired concentrations in M9 buffer. 10 µl drops of serotonin, CNO, or a combination of the two were placed on the lid of a 96-well plate. At time 0 a single worm was placed in each drop of drugged buffer and after 30 minutes the number of eggs by each worm laid was counted under a dissecting microscope.

### Optogenetic activation of photoactivatable adenylyl cyclase to induce egg laying

Animals were staged as late L4 larvae for assay 24 hours later. A photoactivatable adenylyl cyclase (PAC) from *Beggiatoa sp* (amplified from pET28a-ec_bPAC, a gift from Peter Hegemann (Addgene plasmid # 28135)) or empty vector control was transgenically expressed in the egg-laying muscles of *C. elegans* with a *lite-1 (ce314)* background. Worms were kept in foil covered boxes and maintained quickly under dim light to avoid premature activation of the PAC. 24 hours prior to the experiment, single L4 worms were transferred to a new NGM plate containing OP50 and returned to the dark. Both the PAC and empty vector control used the *unc-103e* promoter and were co-injected with *unc-103ep::mCherry*. On the day of the experiment only animals with visible mCherry in their vulva muscles were selected to be assayed. A Leica M165FC microscope equipped with GFP filter set and a digital camera was used to record the experiment. The camera’s exposure settings were adjusted so the activation of the GFP filters set’s blue emission light would be visible on screen. At time 0 the worm was illuminated with 18.2 mW/cm^2^ of 470 ± 20 nm blue light from the microscope’s GFP filter set. The number of eggs laid during 1 minute of blue light illumination was recorded.

### Quantification of unlaid eggs

Animals were staged as L4 larva 30 hours prior to assay. Quantitation of unlaid eggs was performed as described in Chase and Koelle (2004).

### Statistical analysis

Error bars shown in all graphs represent 95% confidence intervals. All statistical analysis was analyzed using GraphPad Prism version 9.3.1 software. Calcium imaging transients in the vm1 and vm2 muscles (Figure 3) were analyzed using a contingency analysis and Fisher’s exact test with two-sided P-values and Bonferroni correction method for multiple comparisons. Egg-laying assays involving CNO-induced activation of the DREADD Gα_q_ receptor were analyzed in Figure 7 using the unpaired t-test with a two-tailed P value with the assumption that both populations had the same standard deviation and in Figure 8 using two-way ANOVA analysis with Šídák’s multiple comparisons test. Egg-laying assays involving optogenetic activation of photoactivatable adenylyl cyclase were analyzed unpaired t-test with a two-tailed P value and assumed both populations had the same standard deviation. All other statistical analyses were performed using one-way ANOVA analysis with Šídák’s multiple comparisons test.

## Data and software availability

## Acknowledgements

Some strains were provided by the CGC, which is funded by NIH Office of Research Infrastructure Programs (P40 OD010440) and the Mitani laboratory, Japan National BioResource Project (NBRP). This work was funded by NIH R01 grants NS086932 and NS036918. We thank the Yale Center for Advanced Light Microscopy Facility for their assistance with confocal microscopy.

## Author Contributions

Andrew C. Olson:

conceptualization, methodology, validation, formal analysis, investigation, resources, writing-original draft, writing review & editing, visualization, supervision, project administration

Allison M. Butt:

validation, investigation, writing review & editing

Nakeirah T.M. Christie:

investigation, writing review & editing, visualization

Ashish Shelar:

Software

Michael R. Koelle:

conceptualization, methodology, writing-original draft, writing review & editing, visualization, supervision, project administration, funding acquisition

---

**Figure 1-figure supplement 1.**
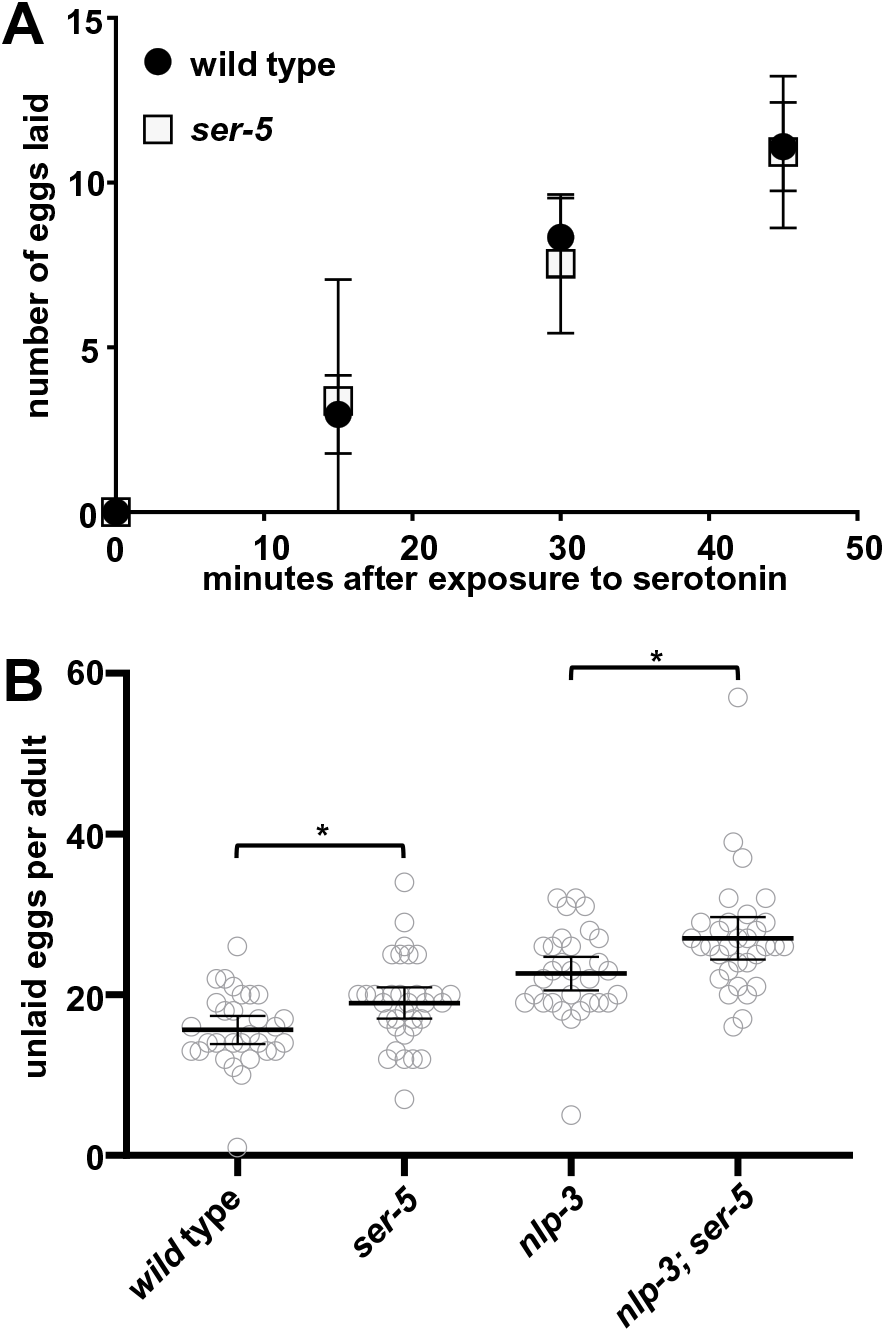
The SER-5 serotonin receptor has only minor effects on egg laying. (**A**) Results of a time course in which worms were placed on plates containing 26 mM serotonin. The number of eggs laid was counted for wild-type and *ser-5* mutant animals. The assay was repeated with 10 worms/plate at least three times per genotype. (**B**) Average number of unlaid eggs per adult worm, n≥30 for each genotype. * = p<0.05. All measurements are given with 95% confidence intervals.

**Figure 1-figure supplement 2.**
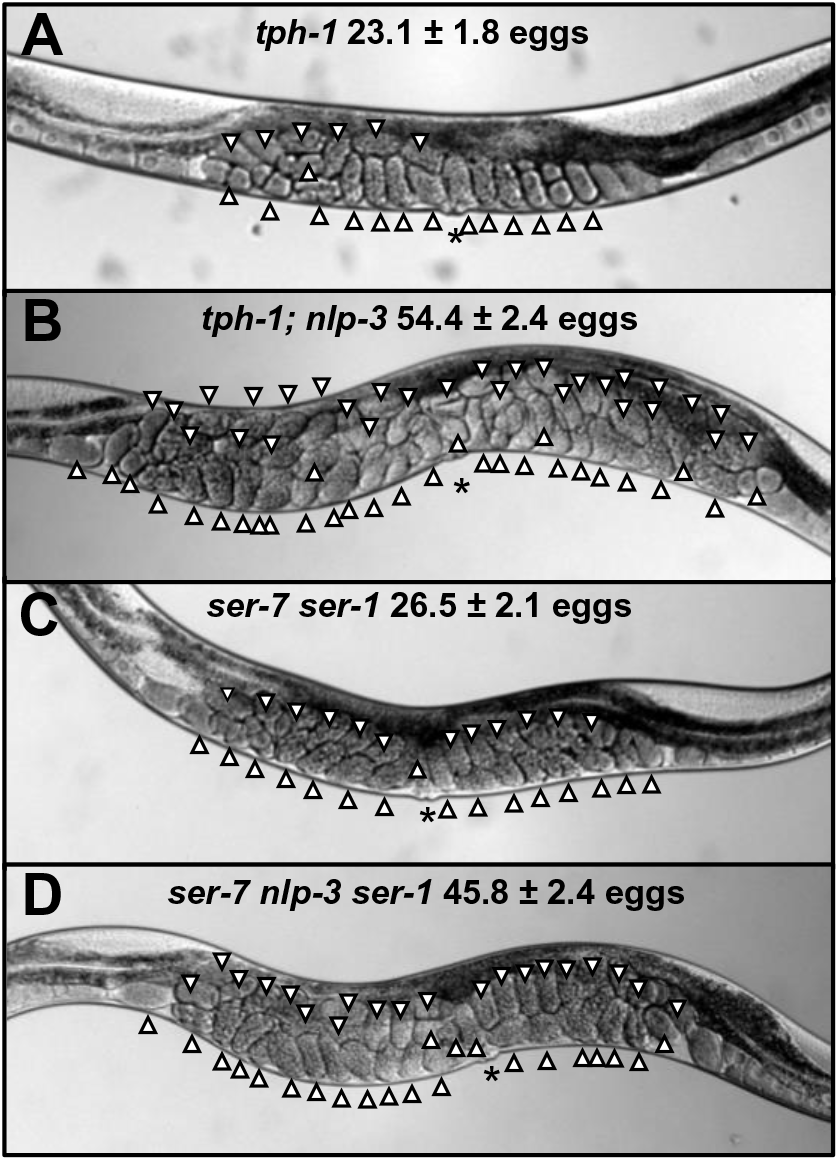
Egg accumulation in additional genotypes. (**A-D**) Photographs of worms of the indicated genotypes, with unlaid eggs indicated by arrowheads. The vulval slit is indicated by *. The average number of unlaid eggs for each genotype is indicated. All measurements are given with 95% confidence intervals.

**Figure 2-figure supplement 1.**
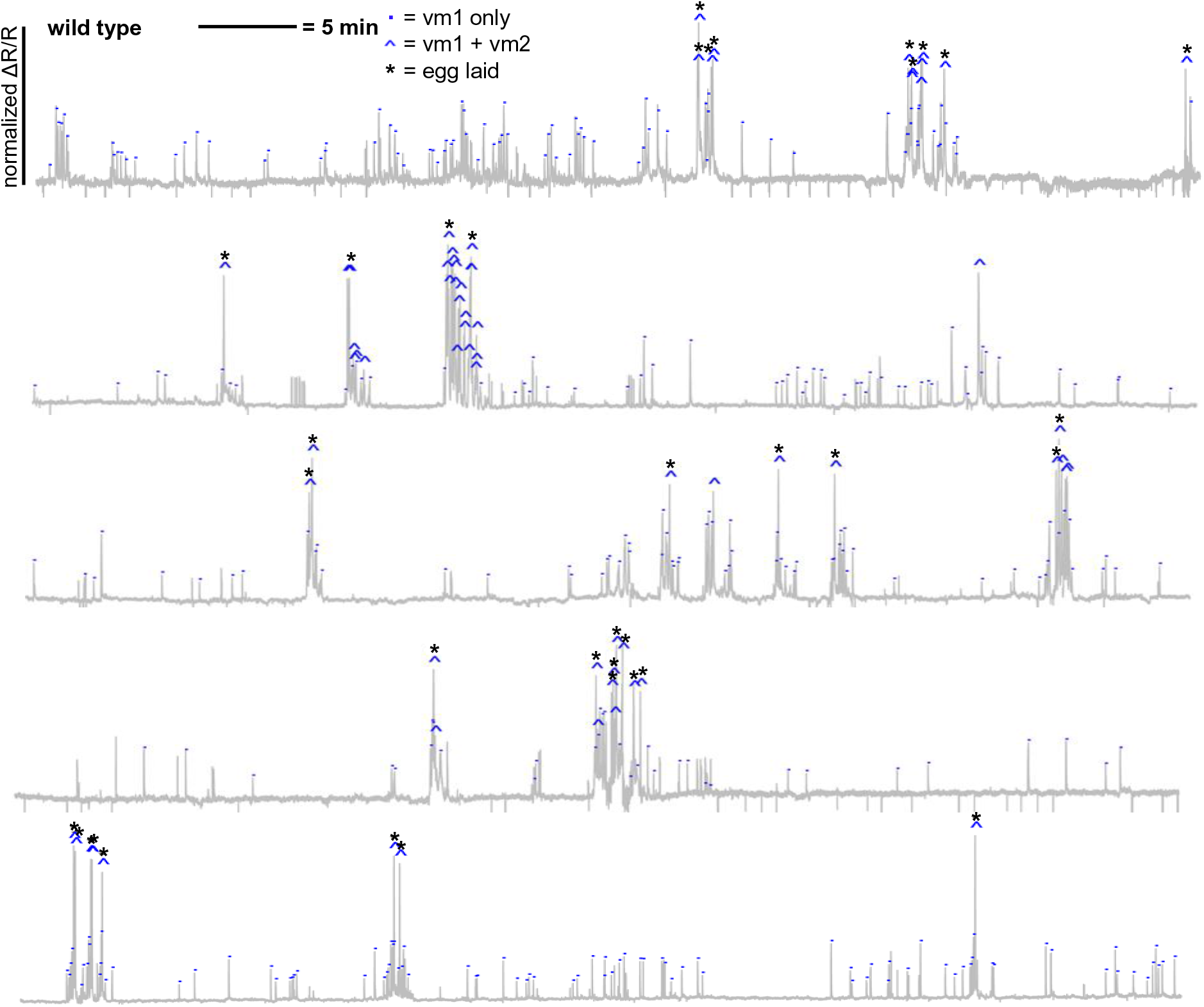
Vulval muscle Ca^2+^ traces in a wild-type background, recorded for one hour each in five different animals.

**Figure 2-figure supplement 2.**
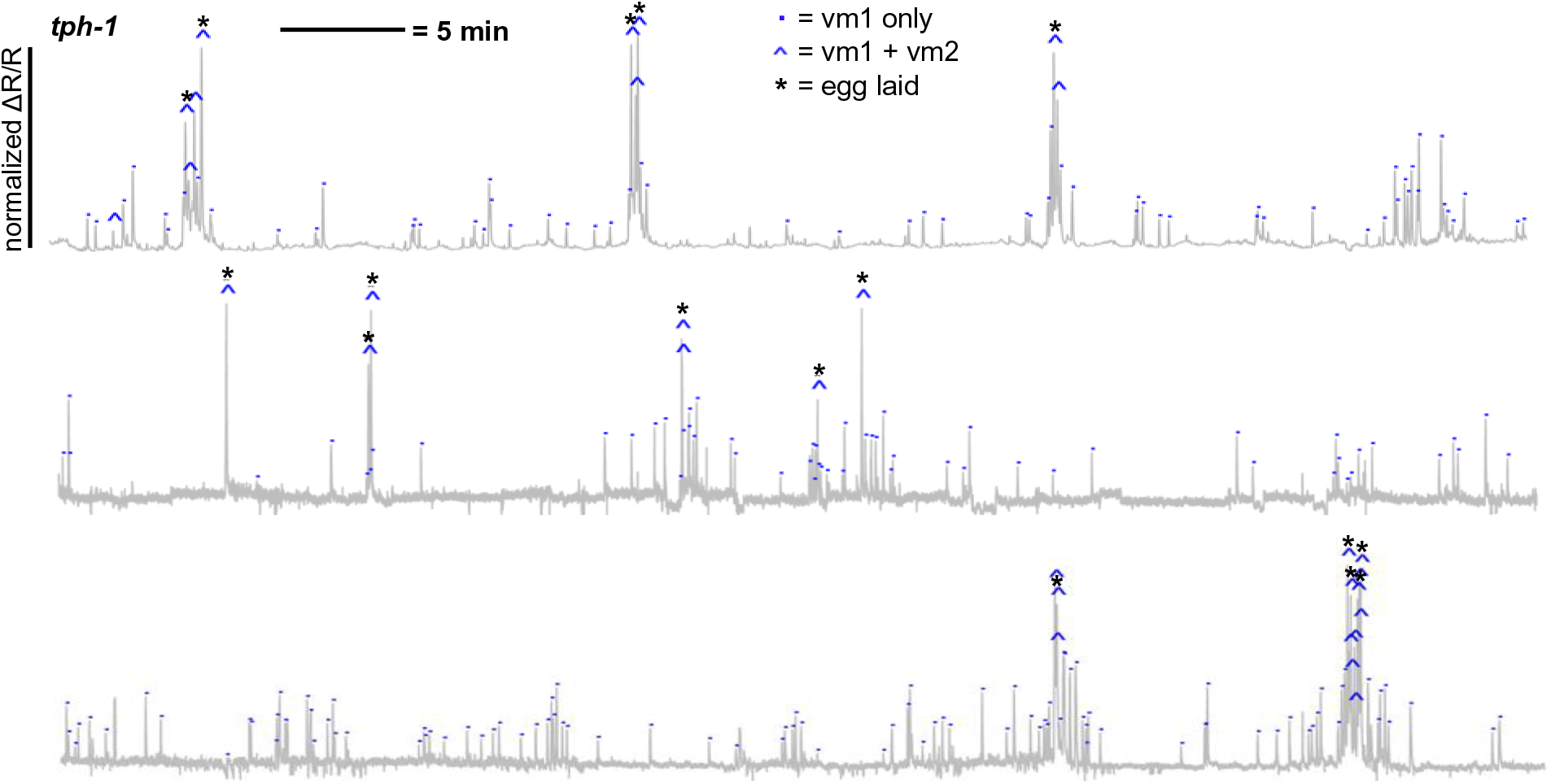
Vulval muscle Ca^2+^ traces in a *tph-1* null mutant background, recorded for one hour each in three different animals.

**Figure 2-figure supplement 3.**
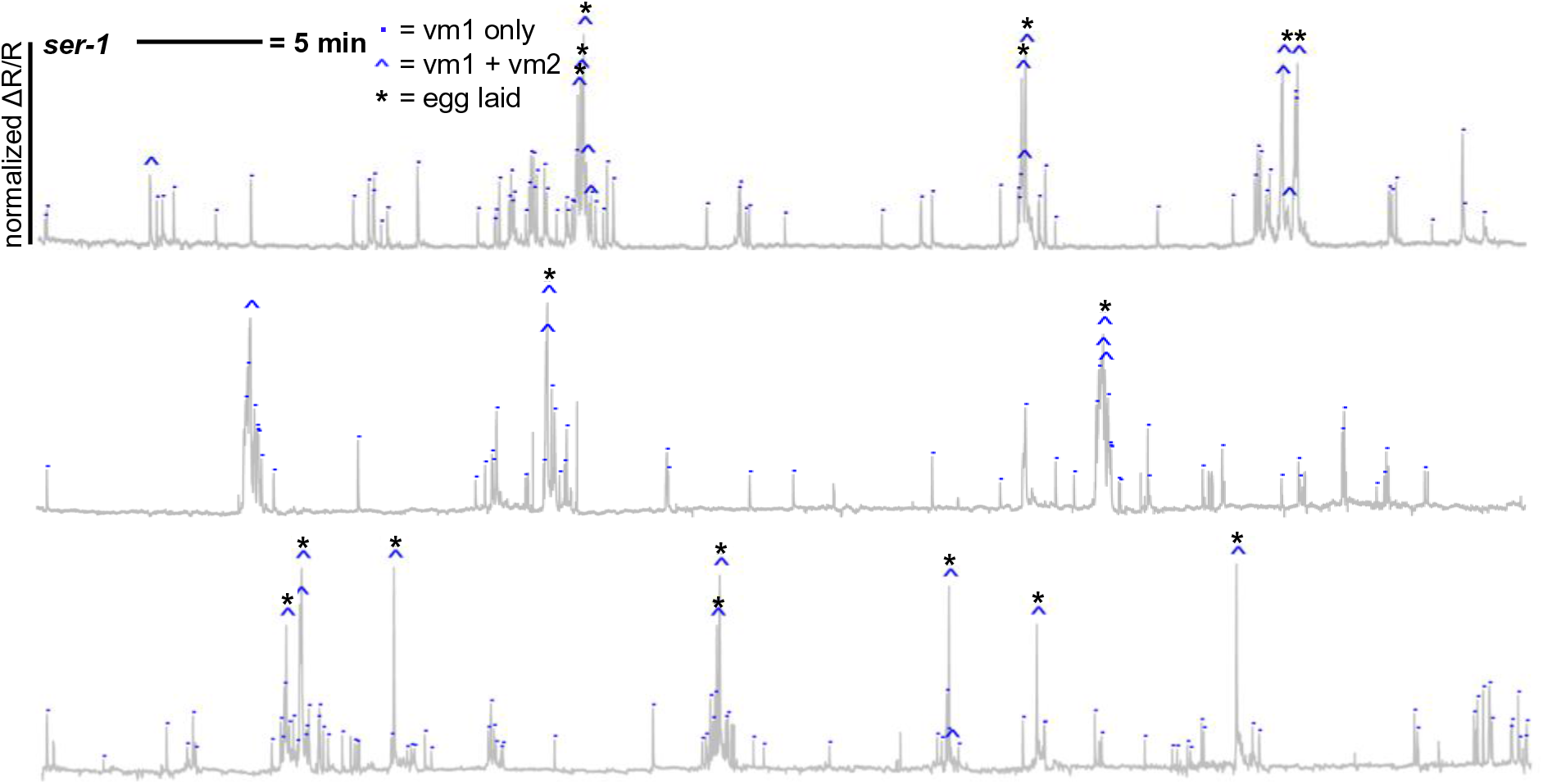
Vulval muscle Ca^2+^ traces in a *ser-1* null mutant background, recorded for one hour each in three different animals.

**Figure 2-figure supplement 4.**
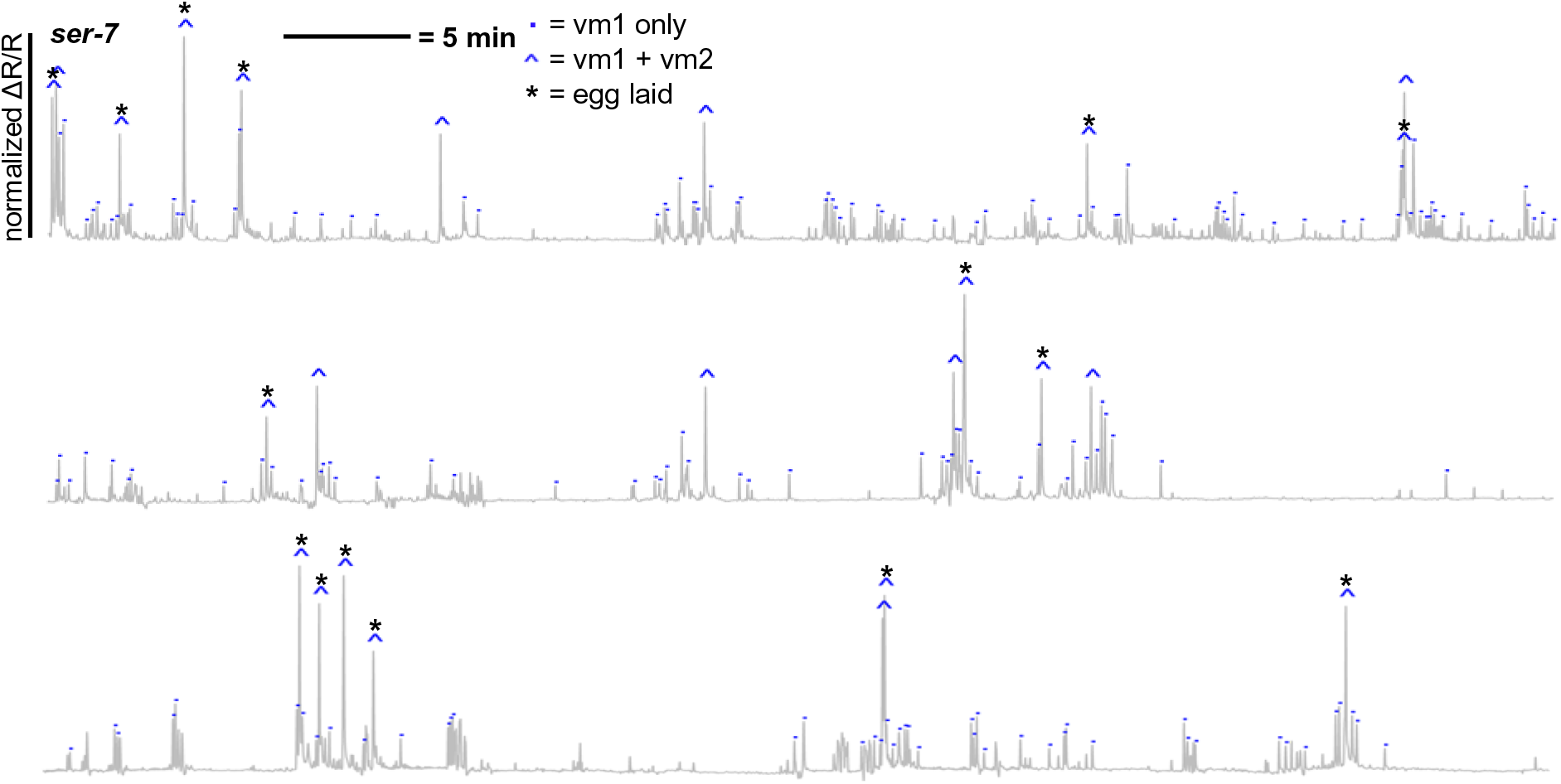
Vulval muscle Ca^2+^ traces in a *ser-7* null mutant background, recorded for one hour each in three different animals.

**Figure 2-figure supplement 5.**
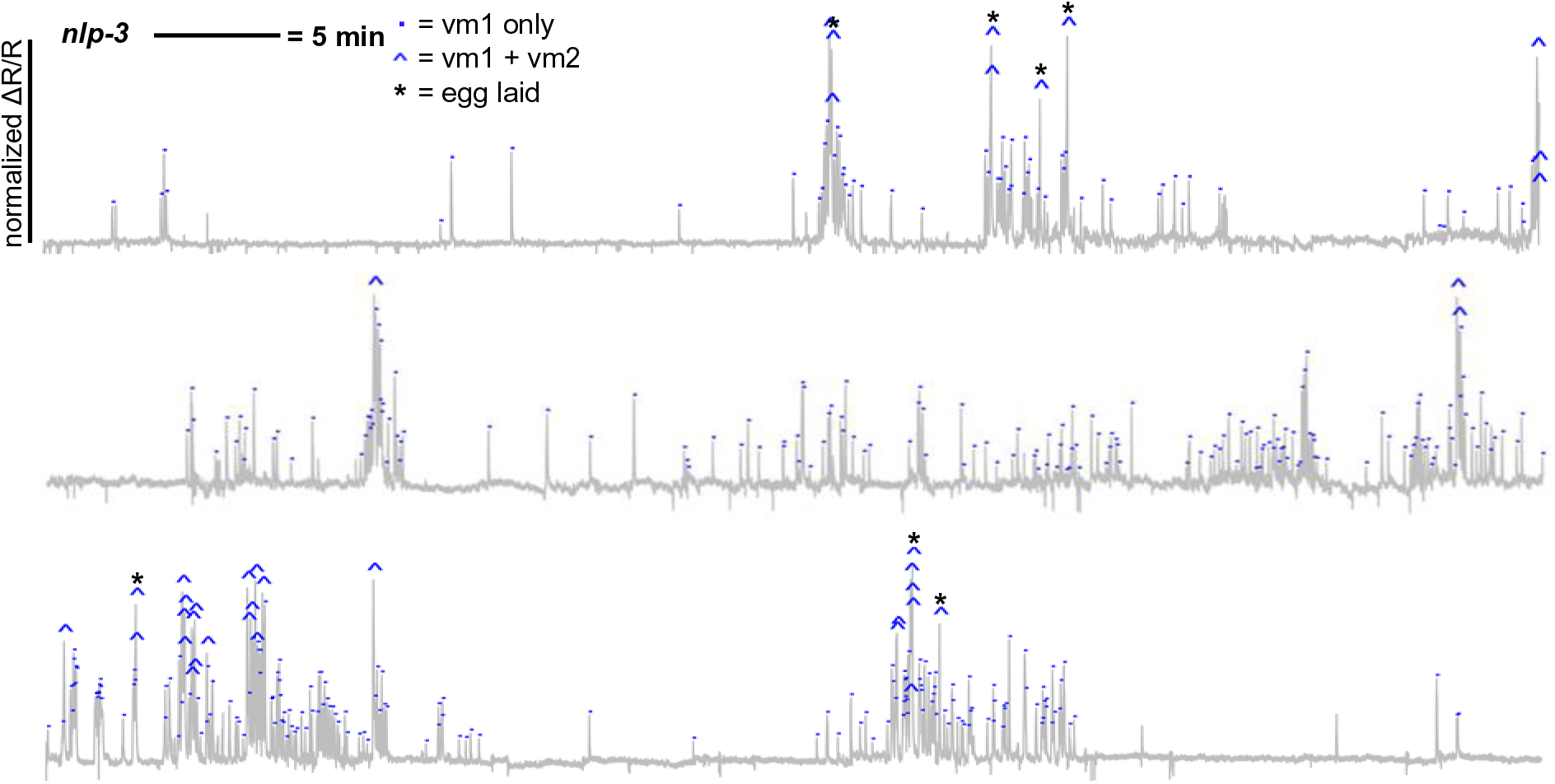
Vulval muscle Ca^2+^ traces in a *nlp-3* null mutant background, recorded for one hour each in three different animals.

**Figure 2-figure supplement 6.**
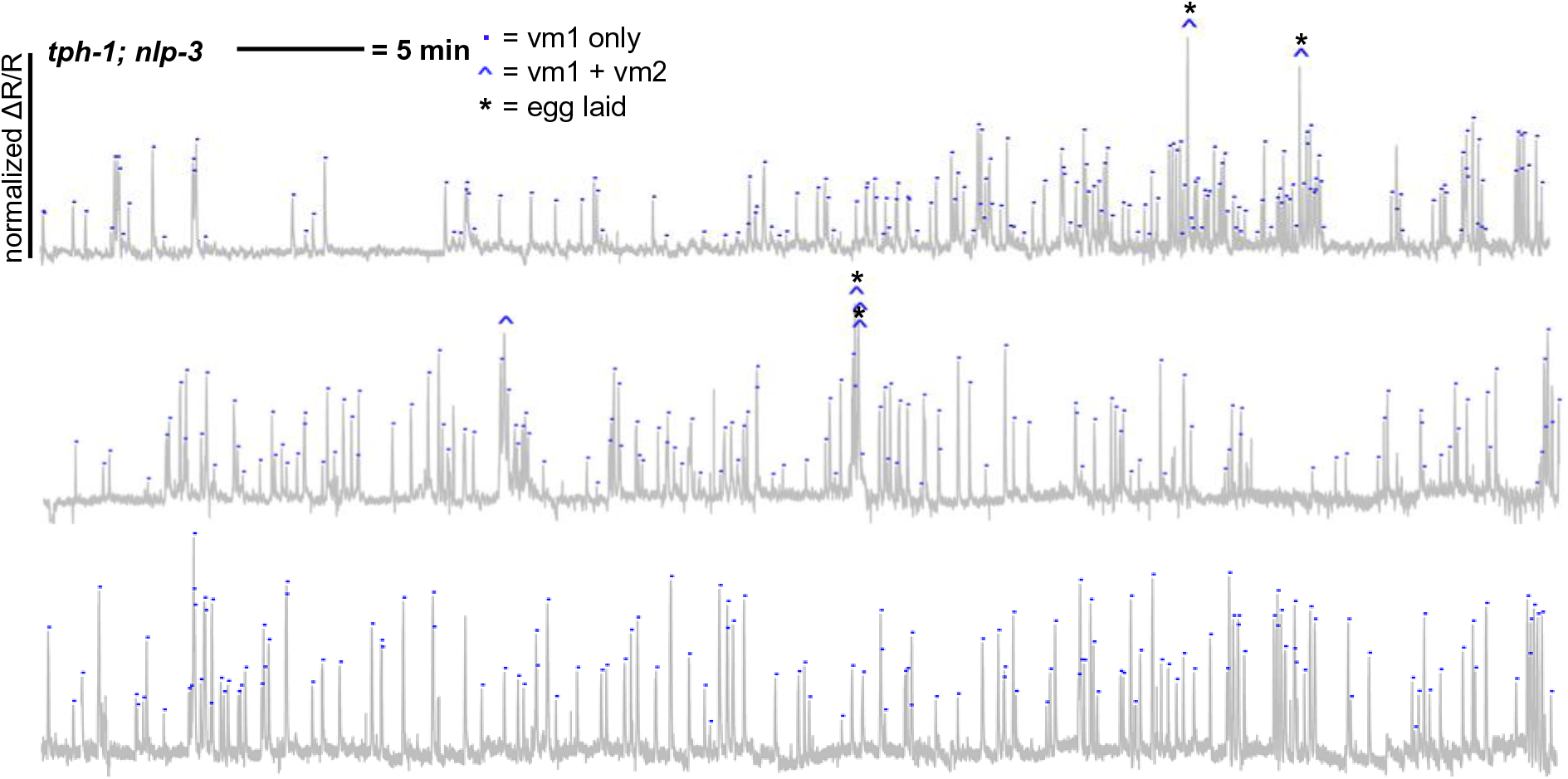
Vulval muscle Ca^2+^ traces in a *tph-1; nlp-3* double null mutant background, recorded for one hour each in three different animals.

**Figure 2-figure supplement 7.**
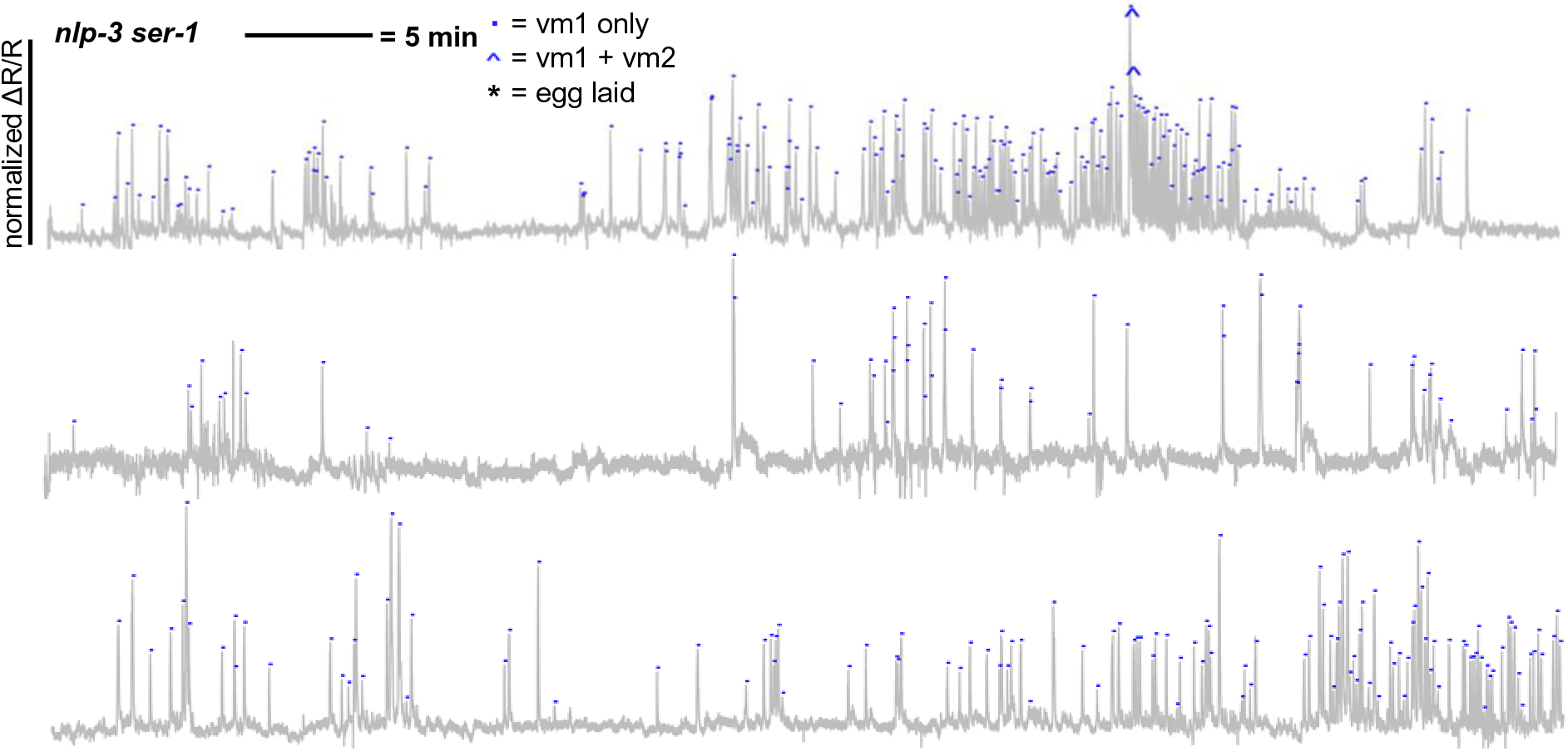
Vulval muscle Ca^2+^ traces in a *nlp-3 ser-1* double null mutant background, recorded for one hour each in three different animals.

**Figure 2-figure supplement 8.**
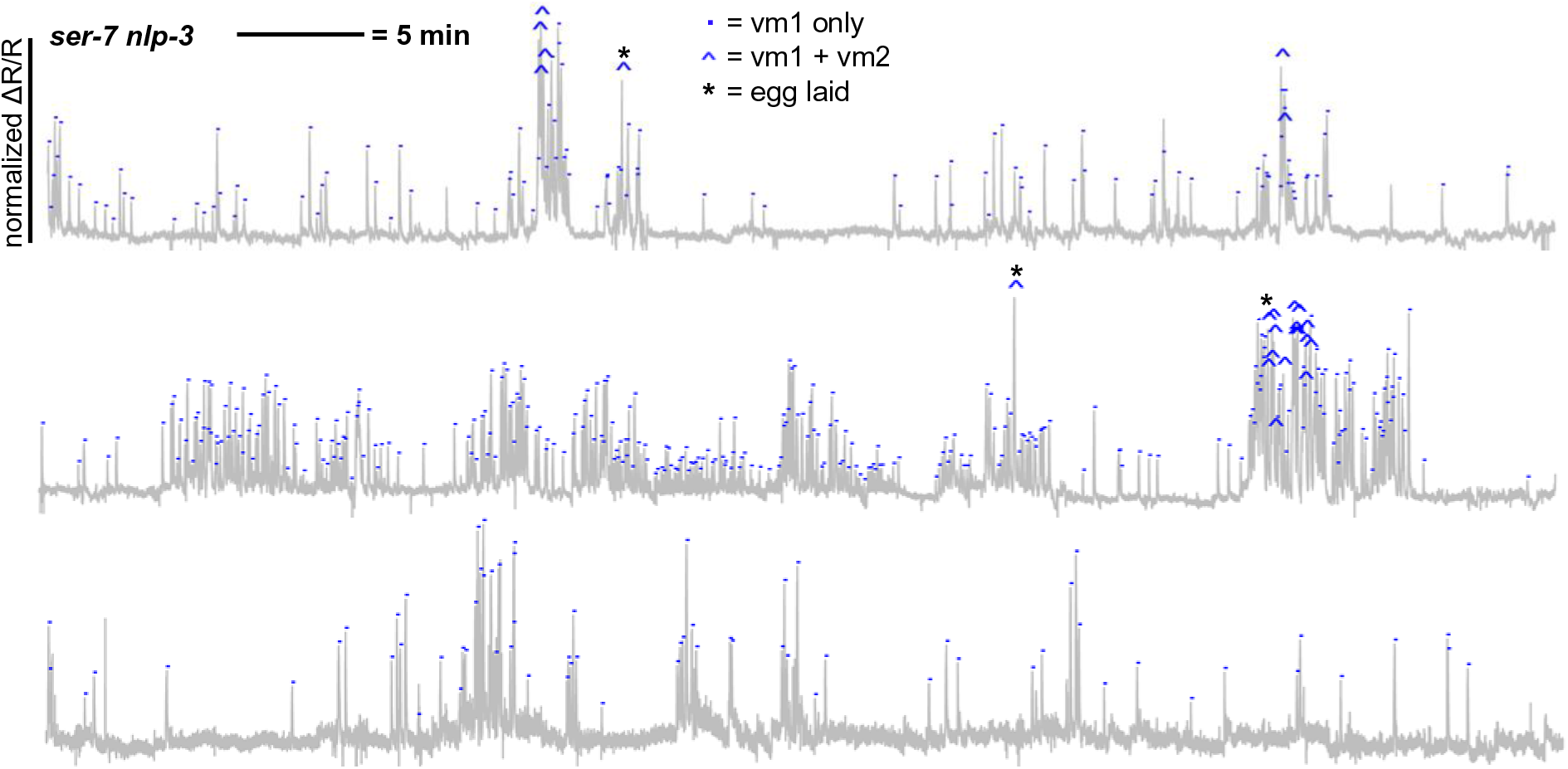
Vulval muscle Ca^2+^ traces in a *ser-7 nlp-3* double null mutant background, recorded for one hour each in three different animals.

**Figure 4-figure supplement 1.**
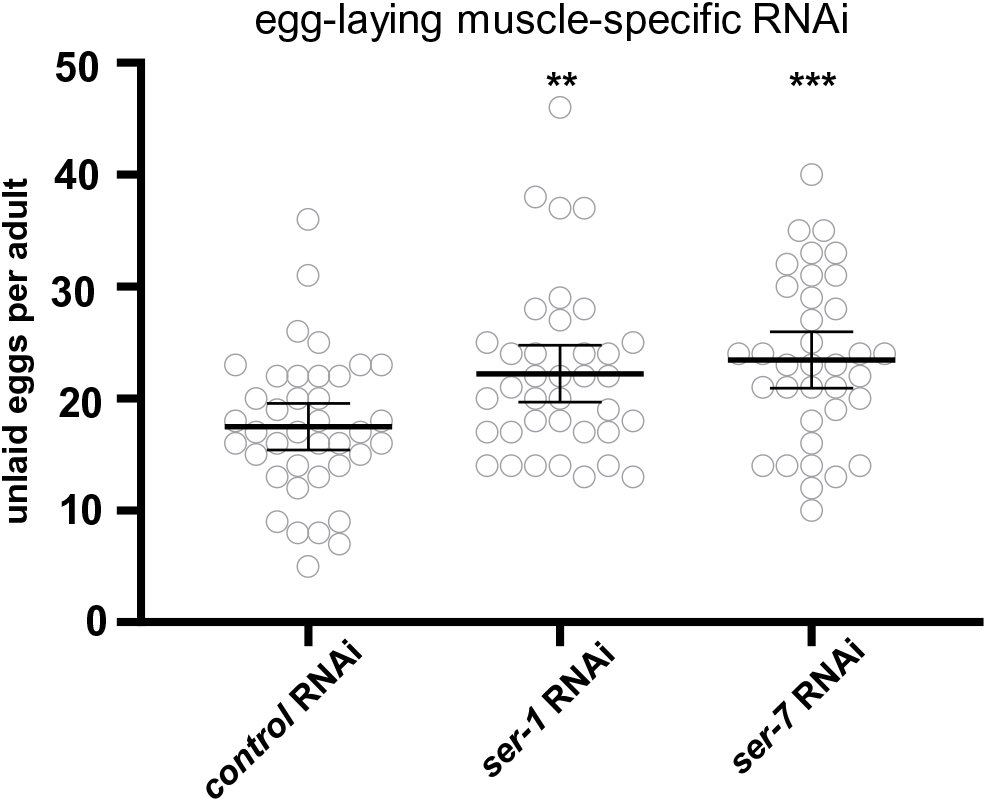
Egg-laying muscle specific knockdown of *ser-1* or *ser-7* result in significantly increased accumulation of unlaid eggs. Egg-laying muscle specific RNAi was used to knock down serotonin receptors and the resulting accumulation of unlaid eggs was measured. Anti-*gfp* RNAi was used as a negative control. Graph indicates the average number of unlaid eggs per adult worm, n≥30 for each genotype. ** = p<0.01, *** = p<0.001. All measurements are given with 95% confidence intervals.

**Figure 5-figure supplement 1.**
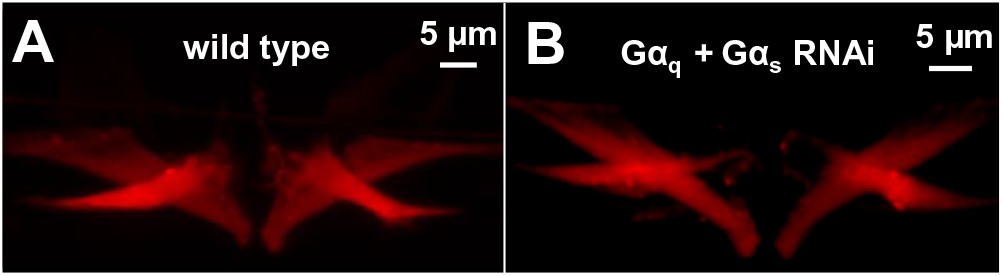
Knock down of Gα_q_ and Gα_s_ in the egg-laying muscles does not disrupt development of the vm1 and vm2 vulval muscle cells. Confocal images of mCherry-labelled vm1 and vm2 vulval muscles in (**A**) adult wild-type worms and (**B**) adult worms in which Gα_q_ and Gα_s_ were both knocked down in the egg-laying muscles. No significant differences are discernable in the morphology of the muscles in these two types of animals. Fifteen animals of each genotype were inspected and the vm1 and mv2 vulval muscles had fully developed in all of the inspected animals.

**Figure 6-figure supplement 1.**
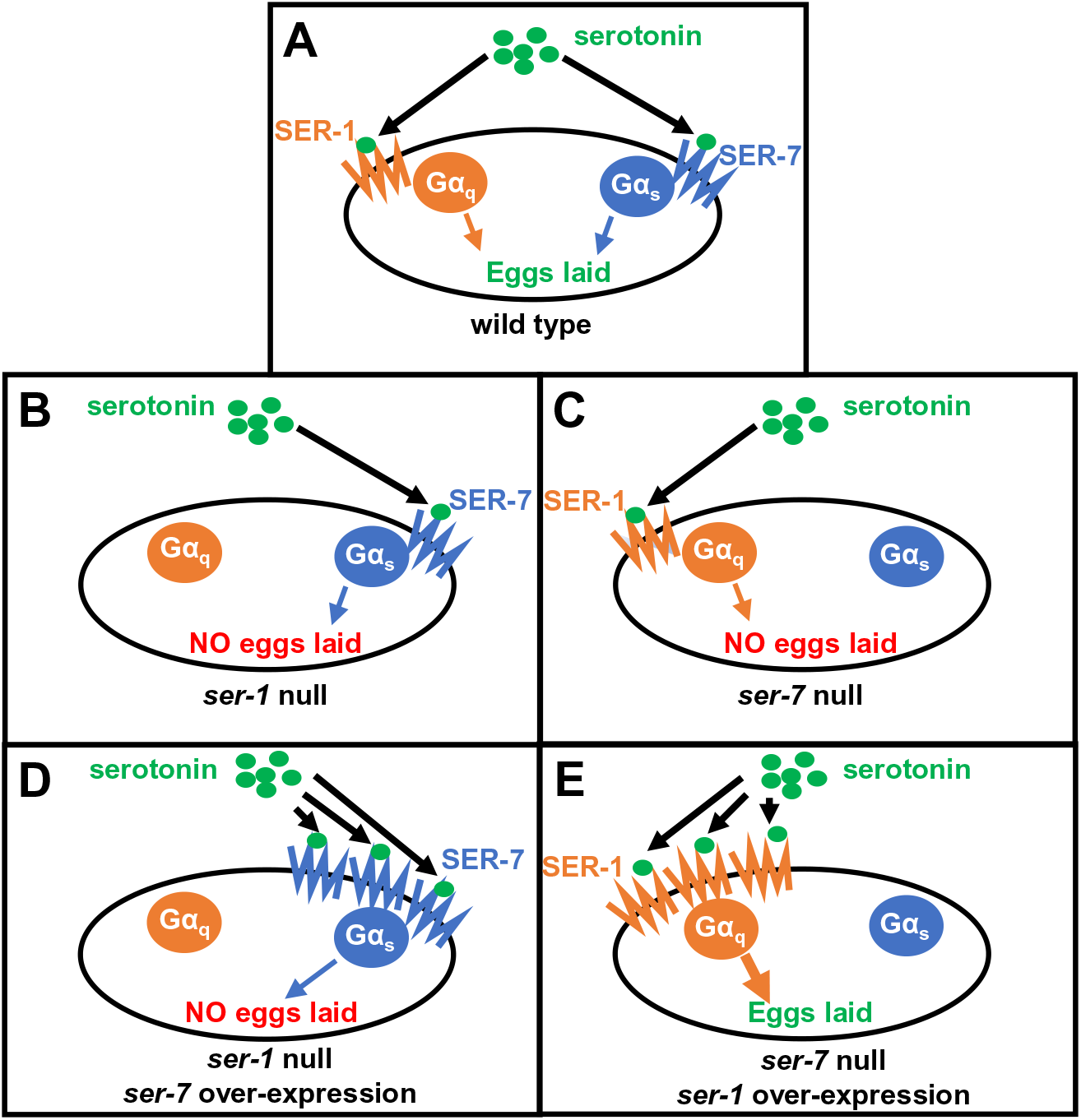
Schematic of the design and results of an experiment overexpressing SER-1 or SER-7 in the absence of the other receptor. (**A**) In wild-type animals, serotonin signals through endogenous levels of SER-1 and SER-7 receptors to activate low levels of Gα_q_ and Gα_s_ signaling, respectively, that combine to induce egg laying. (**B**) Signaling by endogenous SER-7/Gα_s_ is not sufficient to induce egg laying in the absence of SER-1. (**C**) Signaling by endogenous SER-1/Gα_q_ is not sufficient to induce egg laying in the absence of SER-7. (**D**) Overexpressed SER-7 receptor, which might be expected to increase Gα_s_ signaling, is also not sufficient to allow serotonin induce egg laying in the absence of SER-1. (**E**) Overexpressed SER-1 receptor, which might be expected to increase Gα_q_ signaling, is sufficient to allow serotonin to induce egg laying in the absence of SER-7.

**Video 1. Optogenetic activation of Photoactivatable Adenylyl Cyclase (PAC) in the *C. elegans* egg-laying muscles is sufficient to induce egg laying.** The experiment begins with a transgenic worm, expressing PAC in its egg-laying muscles, crawling on a standard laboratory nematode growth medium plate backlit by dim white light. Three seconds into the video, blue light illuminates the worm. Five seconds after blue light illumination the worm lays its first egg. Six eggs are laid within 29 seconds of blue light illumination.

## Supplemental File 1

An excel file containing descriptions of the strains, transgenes, RNAi constructs, and plasmids used in this study.

## References

Bany IA, Dong MQ, Koelle MR. 2003. Genetic and cellular basis for acetylcholine inhibition of *Caenorhabditis elegans* egg-laying behavior. J Neurosci. 23:8060–8069. doi: 10.1523/JNEUROSCI.23-22-08060.2003. PMID: 12954868.

Bargmann CI, Hartwieg E, Horvitz HR. 1993. Odorant-selective genes and neurons mediate olfaction in *C. elegans*. Cell 74:515–527. doi: 10.1016/0092-8674(93)80053-h. PMID: 8348618.

Bonn M, Schmitt A, Lesch KP, Van Bockstaele EJ, Asan E. 2013. Serotonergic innervation and serotonin receptor expression of NPY-producing neurons in the rat lateral and basolateral amygdaloid nuclei. Brain Struct Funct. 218:421–435. doi: 10.1007/s00429-012-0406-5. PMID: 22527118.

Brewer JC, Olson AC, Collins KM, Koelle MR. 2019. Serotonin and neuropeptides are both released by the HSN command neuron to initiate *Caenorhabditis elegans* egg laying. PLoS Genet. 15:e1007896. doi: 10.1371/journal.pgen.1007896. PMID: 30677018.

Bristow MR, Ginsburg R, Umans V, Fowler M, Minobe W, Rasmussen R, Zera P, Menlove R, Shah P, Jamieson S. 1986. Beta 1- and beta 2-adrenergic-receptor subpopulations in nonfailing and failing human ventricular myocardium: coupling of both receptor subtypes to muscle contraction and selective beta 1-receptor down-regulation in heart failure. Circ Res. 59:297–309. doi: 10.1161/01.res.59.3.297. PMID: 2876788.

Carnell L, Illi J, Hong SW, McIntire SL. 2005. The G-protein-coupled serotonin receptor SER-1 regulates egg laying and male mating behaviors in *Ca*e*norhabditis elegans*. J Neurosci. 25:10671–10681. doi: 10.1523/JNEUROSCI.3399-05.2005. PMID: 16291940.

Carre-Pierrat M, Baillie D, Johnsen R, Hyde R, Hart A, Granger L, Ségalat L. 2006. Characterization of the *Caenorhabditis elegans* G protein-coupled serotonin receptors. Invert Neurosci. 6:189–205. doi: 10.1007/s10158-006-0033-z. PMID: 17082916.

Chase DL, Koelle MR. 2004. Genetic analysis of RGS protein function in *Caenorhabditis elegans*. Methods Enzymol. 389:305–320. doi: 10.1016/S0076-6879(04)89018-9. PMID: 15313573.

Chikumi H, Vázquez-Prado J, Servitja JM, Miyazaki H, Gutkind JS. 2002. Potent activation of RhoA by Galpha q and Gq-coupled receptors. J Biol Chem. 277:27130–27134. doi: 10.1074/jbc.M204715200. PMID: 12016230.

Collins KM, Koelle MR. 2013. Postsynaptic ERG potassium channels limit muscle excitability to allow distinct egg-laying behavior states in Caenorhabditis elegans. J Neurosci. 33:761–775. doi: 10.1523/JNEUROSCI.3896-12.2013. PMID: 23303953.

Collins KM, Bode A, Fernandez RW, Tanis JE, Brewer JC, Creamer MS, Koelle MR. 2016. Activity of the *C. elegans* egg-laying behavior circuit is controlled by competing activation and feedback inhibition. Elife 5:e21126. doi: 10.7554/eLife.21126. PMID: 27849154.

Dempsey CM, Mackenzie SM, Gargus A, Blanco G, Sze JY. 2005. Serotonin (5HT), fluoxetine, imipramine and dopamine target distinct 5HT receptor signaling to modulate *Caenorhabditis elegans* egg-laying behavior. Genetics 169:1425–1436. doi: 10.1534/genetics.104.032540. Epub 2005 Jan 16. PMID: 15654117.

Desai C, Horvitz HR. 1989. *Caenorhabditis elegans* mutants defective in the functioning of the motor neurons responsible for egg laying. Genetics 121:703–721. doi: 10.1093/genetics/121.4.703. PMID: 2721931.

Dhakal P, Chaudhry SI, Signorelli R, Collins KM. 2022. Serotonin signals through postsynaptic Gαq, Trio RhoGEF, and diacylglycerol to promote *Caenorhabditis elegans* egg-laying circuit activity and behavior. Genetics 221:iyac084. doi: 10.1093/genetics/iyac084. PMID: 35579369.

Esposito G, Di Schiavi E, Bergamasco C, Bazzicalupo P. 2007. Efficient and cell specific knock-down of gene function in targeted *C. elegans* neurons. Gene 395:170–176. doi: 10.1016/j.gene.2007.03.002. PMID: 17459615.

Feng J, Cai X, Zhao J, Yan Z. 2001. Serotonin receptors modulate GABA(A) receptor channels through activation of anchored protein kinase C in prefrontal cortical neurons. J Neurosci. 21:6502–6511. doi: 10.1523/JNEUROSCI.21-17-06502.2001. PMID: 11517239.

Fernandez RW, Wei K, Wang EY, Mikalauskaite D, Olson A, Pepper J, Christie N, Kim S, Weissenborn S, Sarov M, Koelle MR. 2020. Cellular Expression and Functional Roles of All 26 Neurotransmitter GPCRs in the *C. elegans* Egg-Laying Circuit. J Neurosci. 40:7475–7488. doi: 10.1523/JNEUROSCI.1357-20.2020. PMID: 32847964.

Hanlon CD, Andrew DJ. 2015. Outside-in signaling--a brief review of GPCR signaling with a focus on the Drosophila GPCR family. J Cell Sci. 128:3533–35342. doi: 10.1242/jcs.175158. PMID: 26345366.

Hamdan FF, Ungrin MD, Abramovitz M, Ribeiro P. 1999. Characterization of a novel serotonin receptor from *Caenorhabditis elegans*: cloning and expression of two splice variants. J Neurochem. 72:1372–1383. doi: 10.1046/j.1471-4159.1999.721372.x. PMID: 10098838.

Hapiak VM, Hobson RJ, Hughes L, Smith K, Harris G, Condon C, Komuniecki P, Komuniecki RW. 2009. Dual excitatory and inhibitory serotonergic inputs modulate egg laying in *Caenorhabditis elegans*. Genetics 181:153–163. doi: 10.1534/genetics.108.096891. PMID: 19001289.

Harris G, Mills H, Wragg R, Hapiak V, Castelletto M, Korchnak A, Komuniecki RW. 2010. The monoaminergic modulation of sensory-mediated aversive responses in *Caenorhabditis elegans* requires glutamatergic/peptidergic cotransmission. J Neurosci. 30:7889–7899. doi: 10.1523/JNEUROSCI.0497-10.2010. PMID: 20534837.

Henss T, Schneider M, Vettkötter D, Costa WS, Liewald JF, Gottschalk A. 2022. Photoactivated Adenylyl Cyclases as Optogenetic Modulators of Neuronal Activity. Methods Mol Biol. 2483:61–76. doi: 10.1007/978-1-0716-2245-2_4. PMID: 35286669.

Hobson RJ, Geng J, Gray AD, Komuniecki RW. 2003. SER-7b, a constitutively active Galphas coupled 5-HT7-like receptor expressed in the *Caenorhabditis elegans* M4 pharyngeal motorneuron. J Neurochem. 87:22–29. doi: 10.1046/j.1471-4159.2003.01967.x. PMID: 12969249.

Hobson RJ, Hapiak VM, Xiao H, Buehrer KL, Komuniecki PR, Komuniecki RW. 2006. SER-7, a *Caenorhabditis elegans* 5-HT7-like receptor, is essential for the 5-HT stimulation of pharyngeal pumping and egg laying. Genetics 172:159–169. doi: 10.1534/genetics.105.044495. PMID: 16204223.

Jiang H, Galtes D, Wang J, Rockman HA. 2022. G Protein-Coupled Receptor Signaling: Transducers and Effectors. Am J Physiol Cell Physiol. doi: 10.1152/ajpcell.00210.2022. PMID: 35816644.

Kaur H, Carvalho J, Looso M, Singh P, Chennupati R, Preussner J, Günther S, Albarrán-Juárez J, Tischner D, Classen S, Offermanns S, Wettschureck N. 2017. Single-cell profiling reveals heterogeneity and functional patterning of GPCR expression in the vascular system. Nat Commun. 8:15700. doi: 10.1038/ncomms15700. 10:1448. PMID: 28621310.

Kopchock RJ 3rd, Ravi B, Bode A, Collins KM. 2021. The Sex-Specific VC Neurons Are Mechanically Activated Motor Neurons That Facilitate Serotonin-Induced Egg Laying in *C. elegans*. J Neurosci. 41:3635–3650. doi: 10.1523/JNEUROSCI.2150-20.2021. PMID: 33687965.

Lee HM, Giguere PM, Roth BL. 2014. DREADDs: novel tools for drug discovery and development. Drug Discov Today. 19:469–473. doi:10.1016/j.drudis.2013.10.018.

Lin F, Owens WA, Chen S, Stevens ME, Kesteven S, Arthur JF, Woodcock EA, Feneley MP, Graham RM. 2001. Targeted alpha(1A)-adrenergic receptor overexpression induces enhanced cardiac contractility but not hypertrophy. Circ Res. 89:343–350. doi: 10.1161/hh1601.095912. PMID: 11509451.

Lopez ER, Carbajal AG, Tian JB, Bavencoffe A, Zhu MX, Dessauer CW, Walters ET. 2021. Serotonin enhances depolarizing spontaneous fluctuations, excitability, and ongoing activity in isolated rat DRG neurons via 5-HT_4_ receptors and cAMP-dependent mechanisms. Neuropharmacology. 184:108408. doi: 10.1016/j.neuropharm.2020.108408. PMID: 33220305.

Lutz S, Freichel-Blomquist A, Yang Y, Rümenapp U, Jakobs KH, Schmidt M, Wieland T. 2005. The guanine nucleotide exchange factor p63RhoGEF, a specific link between Gq/11-coupled receptor signaling and RhoA. J Biol Chem. 280:11134–11139. doi: 10.1074/jbc.M411322200. PMID: 15632174.

Lutz S, Shankaranarayanan A, Coco C, Ridilla M, Nance MR, Vettel C, Baltus D, Evelyn CR, Neubig RR, Wieland T, Tesmer JJ. 2007. Structure of Galphaq-p63RhoGEF-RhoA complex reveals a pathway for the activation of RhoA by GPCRs. Science 318:1923–7. doi: 10.1126/science.1147554. PMID: 18096806.

Lymperopoulos A, Cora N, Maning J, Brill AR, Sizova A. 2021. Signaling and function of cardiac autonomic nervous system receptors: Insights from the GPCR signalling universe. FEBS J. 288:2645–2659. doi: 10.1111/febs.15771. PMID: 33599081.

Marder E, Bucher D. 2007. Understanding circuit dynamics using the stomatogastric nervous system of lobsters and crabs. Annu Rev Physiol. 69:291–316. doi: 10.1146/annurev.physiol.69.031905.161516. PMID: 17009928.

McCloskey DT, Turnbull L, Swigart P, O’Connell TD, Simpson PC, Baker AJ. 2003. Abnormal myocardial contraction in alpha(1A)- and alpha(1B)-adrenoceptor double-knockout mice. J Mol Cell Cardiol. 35:1207–1216. doi: 10.1016/s0022-2828(03)00227-x. PMID: 14519431.

Meggs LG, Coupet J, Huang H, Cheng W, Li P, Capasso JM, Homcy CJ, Anversa P. 1993. Regulation of angiotensin II receptors on ventricular myocytes after myocardial infarction in rats. Circ Res. 72:1149–1162. doi: 10.1161/01.res.72.6.1149. PMID: 8495545.

Miller KG, Alfonso A, Nguyen M, Crowell JA, Johnson CD, Rand JB. 1996. A genetic selection for *Caenorhabditis elegans* synaptic transmission mutants. Proc Natl Acad Sci U S A. 93:12593–12598. doi: 10.1073/pnas.93.22.12593. PMID: 8901627.

O’Connell TD, Ishizaka S, Nakamura A, Swigart PM, Rodrigo MC, Simpson GL, Cotecchia S, Rokosh DG, Grossman W, Foster E, Simpson PC. 2003. The alpha(1A/C)-and alpha(1B)-adrenergic receptors are required for physiological cardiac hypertrophy in the double-knockout mouse. J Clin Invest. 111:1783–1791. doi: 10.1172/JCI16100. PMID: 12782680.

Prömel S, Fiedler F, Binder C, Winkler J, Schöneberg T, Thor D. 2016. Deciphering and modulating G protein signalling in *C. elegans* using the DREADD technology. Sci Rep. 6:28901. doi: 10.1038/srep28901. PMID: 27461895.

Rasmussen K, Aghajanian GK. 1990. Serotonin excitation of facial motoneurons: receptor subtype characterization. Synapse. 5:324–332. doi: 10.1002/syn.890050409. PMID: 2360199.

Ravi B, Nassar LM, Kopchock RJ 3rd, Dhakal P, Scheetz M, Collins KM. 2018b. Ratiometric Calcium Imaging of Individual Neurons in Behaving *Caenorhabditis Elegans*. J Vis Exp. (**132**):56911. doi: 10.3791/56911. PMID: 29443112.

Reynolds NK, Schade MA, Miller KG. 2005. Convergent, RIC-8-dependent Galpha signaling pathways in the *Caenorhabditis elegans* synaptic signaling network. Genetics 169:651–670. doi: 10.1534/genetics.104.031286. PMID: 15489511.

Ringstad N, Horvitz HR. 2008. FMRFamide neuropeptides and acetylcholine synergistically inhibit egg-laying by *C. elegans*. Nat Neurosci. 11:1168–1176. doi: 10.1038/nn.2186. PMID: 18806786.

Rojas RJ, Yohe ME, Gershburg S, Kawano T, Kozasa T, Sondek J. 2007. Galphaq directly activates p63RhoGEF and Trio via a conserved extension of the Dbl homology-associated pleckstrin homology domain. J Biol Chem. 282:29201–29210. doi: 10.1074/jbc.M703458200. PMID: 17606614.

Salazar NC, Chen J, Rockman HA. 2007. Cardiac GPCRs: GPCR signaling in healthy and failing hearts. Biochim Biophys Acta. 1768:1006–1018. doi: 10.1016/j.bbamem.2007.02.010. PMID: 17376402.

Sarkar P, Mozumder S, Bej A, Mukherjee S, Sengupta J, Chattopadhyay A. 2020. Structure, dynamics and lipid interactions of serotonin receptors: excitements and challenges. Biophys Rev. 13:101–122. doi: 10.1007/s12551-020-00772-8. PMID: 33188638.

Smith SJ, Sümbül U, Graybuck LT, Collman F, Seshamani S, Gala R, Gliko O, Elabbady L, Miller JA, Bakken TE, Rossier J, Yao Z, Lein E, Zeng H, Tasic B, Hawrylycz M. 2019. Single-cell transcriptomic evidence for dense intracortical neuropeptide networks. Elife 8:e47889. doi: 10.7554/eLife.47889. PMID: 31710287.

Smrcka AV, Hepler JR, Brown KO, Sternweis PC. 1991. Regulation of polyphosphoinositide-specific phospholipase C activity by purified Gq. Science 251:804–807. doi: 10.1126/science.1846707. PMID: 1846707.

Steuer Costa W, Yu SC, Liewald JF, Gottschalk A. 2017. Fast cAMP Modulation of Neurotransmission via Neuropeptide Signals and Vesicle Loading. Curr Biol. 27:495–507. doi: 10.1016/j.cub.2016.12.055. PMID: 28162892.

Sze JY, Victor M, Loer C, Shi Y, Ruvkun G. 2000. Food and metabolic signalling defects in a *Caenorhabditis elegans* serotonin-synthesis mutant. Nature 403:560–564. doi: 10.1038/35000609. PMID: 10676966.

Taylor SJ, Chae HZ, Rhee SG, Exton JH. 1991. Activation of the beta 1 isozyme of phospholipase C by alpha subunits of the Gq class of G proteins. Nature 350:516–518. doi: 10.1038/350516a0. PMID: 1707501.

Taylor CW. 2017. Regulation of IP_3_ receptors by cyclic AMP. Cell Calcium 63:48–52. doi: 10.1016/j.ceca.2016.10.005. PMID: 27836216.

Trent C, Tsuing N, Horvitz HR. 1983. Egg-laying defective mutants of the nematode *Caenorhabditis elegans*. Genetics 104:619–647. doi: 10.1093/genetics/104.4.619. PMID: 11813735.

Wang J, Gareri C, Rockman HA. 2018. G-Protein-Coupled Receptors in Heart Disease. Circ Res. 123:716–735. doi: 10.1161/CIRCRESAHA.118.311403. 123:e34. PMID: 30355236.

Waggoner LE, Zhou GT, Schafer RW, Schafer WR. 1998. Control of alternative behavioral states by serotonin in *Caenorhabditis elegans*. Neuron 21:203–214. doi: 10.1016/s0896-6273(00)80527-9. PMID: 9697864.

Williams SL, Lutz S, Charlie NK, Vettel C, Ailion M, Coco C, Tesmer JJ, Jorgensen EM, Wieland T, Miller KG. 2007. Trio’s Rho-specific GEF domain is the missing Galpha q effector in *C. elegans*. Genes Dev. 21:2731–246. doi: 10.1101/gad.1592007. PMID: 17942708.

Winston WM, Molodowitch C, Hunter CP. 2002. Systemic RNAi in *C. elegans* requires the putative transmembrane protein SID-1. Science 295:2456–2459. doi: 10.1126/science.1068836. PMID: 11834782.

Xu H, Shi X, Li X, Zou J, Zhou C, Liu W, Shao H, Chen H, Shi L. 2020. Neurotransmitter and neuropeptide regulation of mast cell function: a systematic review. J Neuroinflammation. 17:356. doi: 10.1186/s12974-020-02029-3. PMID: 33239034.

Xu YJ, Gopalakrishnan V. 1991. Vasopressin increases cytosolic free [Ca2+] in the neonatal rat cardiomyocyte. Evidence for V1 subtype receptors. Circ Res. 69:239–245. doi: 10.1161/01.res.69.1.239. PMID: 2054937.

